# Spatial atlas of gut-draining lymph nodes reveals the developmental organization of intestinal tolerance and immunity

**DOI:** 10.64898/2026.11.16.767093

**Authors:** Yun Lo, Priyanka Lakhiani, Anushka Yadav, Vanja Cabric, Greyson Feather, Ash Binoy, Yoselin A. Paucar Iza, Yollanda Parisotto, Tilman Hoelting, Gayathri Shibu, Tyler Park, Chrysothemis C. Brown

**Author notes:** These authors contributed equally to this work (Yun Lo, Priyanka Lakhiani, Anushka Yadav). Corresponding author. (C.C.B.).

## Abstract

Early life offers a window to establish lasting tolerance to dietary and commensal microbial antigens. How developing gut lymph nodes balance tolerance and immunity remains unclear. Using spatial transcriptomics, we map small intestine- and colon-draining nodes during weaning and adulthood, resolving rare immune and stromal populations and their niches, including Thetis cells (TCs), an antigen-presenting lineage critical for intestinal tolerance. We uncover a peri-weaning subcapsular compartment enriched for TC I but depleted of classical antigen-relaying macrophages, coincident with attenuated follicular responses. Peri-weaning colonic nodes are hubs of T cell activation enriched for peripherally-induced regulatory T (pTreg) cell responses. We resolve distinct classical dendritic cell (cDC) and TC niches, including a pTreg-associated niche shared by TC IV and CCR7⁺ cDC2. We further characterize a distinct population of intestinal tissue TCs, and establish TC migration to draining lymph nodes. Together, our atlas provides a framework for understanding how the timing of intestinal antigen exposure intersects with developing lymph node niches to shape tolerance or immunity.

## INTRODUCTION

The intestinal immune system must balance protective responses to enteric pathogens with tolerance to dietary and commensal microbial antigens. These divergent outcomes are shaped in large part by CD4 T cells, whose differentiation into distinct T helper 1 (T_H_1), T_H_2, T_H_17, T follicular helper (T_FH_) or peripherally-induced regulatory T (pTreg) cells determines the nature of the immune response. Among these, pTreg cells are essential for restraining inflammatory responses to food and microbiota^1–3^. Disruption of this balance contributes to food allergy, inflammatory bowel disease and other immune-mediated disorders^4^.

Early life represents a critical window for the development of intestinal pTreg cells and tolerance^5–8^. The transition from milk to solid food coincides with profound changes in the abundance and composition of the gut microbiota, increased exposure to dietary antigens and a wave of microbiota- and food-specific pTreg cell differentiation^5^. During this period, CD4 T cell responses to intestinal antigens are strongly biased towards tolerance, whereas exposure later in life may elicit alternative effector responses^9–13^. The timing of first exposure can therefore have lasting consequences, as evidenced by the increased risk of food allergy following delayed introduction of allergenic foods^6,14^ . These findings raise a fundamental question: why is the developing intestinal immune system uniquely suited to establish tolerance?

Thetis cells (TCs), comprise a lineage of antigen-presenting cells (APCs), ontogenically distinct from classical dendritic cells (cDCs)^15–18^. Among the four TC subsets, αvββ8^+^ TC IV plays an essential role in promoting microbiota- and food-specific pTreg cell differentiation^9,15–23^. In contrast, cDCs comprising cDC1, cDC2A, cDC2B and their migratory CCR7^+^ counterparts promote distinct effector T cell fates^24^. TCs undergo a wave of differentiation during the peri-weaning period, peaking in number in gut lymph nodes at two weeks of age before declining rapidly thereafter^9,15,25–27^, mirroring the window for enhanced pTreg generation. The developmental balance between cDC and TC populations may therefore contribute to the tolerogenic bias of early-life gut lymph nodes.

TC development is closely linked to lymph node organogenesis and postnatal maturation. TCs share a developmental pathway with lymphoid tissue inducer (LTi) cells^25^, which are essential for lymph node formation, while TC I differentiation depends on RANKL supplied by lymphoid tissue organizer (LTo)-derived marginal reticular cells (MRCs) which are similarly enriched during early life^25^. TC IV are found almost exclusively in gut-draining lymph nodes^9,15^, suggesting that local cues direct their differentiation. Together, these findings implicate the developing lymph node microenvironment in shaping TC abundance and specialization.

However, APC abundance alone may not explain why the same antigen elicits distinct T cell fates at different ages, or why some antigens induce effector responses during early life. Antigen identity, physical form and route of lymph node entry influence which APCs gain access^20,21^. Thus, the division of labor between TC and cDC subsets may depend on both specialized functions and privileged access to particular intestinal antigens. Yet TC heterogeneity has largely been studied in lymph nodes, where these cells are most abundant. Whether TCs acquire dietary and microbial antigens in intestinal tissues before migrating to draining lymph nodes, or capture antigens delivered directly to the lymph node, remains unknown.

Within lymph nodes, the ability of APCs to access antigen depends on their spatial localization. Lymph-borne cell-free antigens enter the subcapsular sinus, where size and physical form influence access to stromal conduits and capture by sinus-associated cells^29^. Subcapsular sinus macrophages (SSMs) relay intact antigen to follicular B cells and follicular dendritic cells (FDCs), supporting B cell-mediated antigen presentation required for terminal T_FH_ differentiation and germinal center responses^30–33^. Resident cDCs sample antigens from lymphatic sinuses and conduits^34–36^, whereas migratory CCR7^+^ cDCs carry intestinal antigens from tissues to the T cell zone for presentation to naïve T cells^34,37,38^.

Much of our understanding of antigen trafficking and APC uptake derives from imaging studies of adult skin-draining lymph nodes, often after infection or immunization. These pathways remain poorly defined in developing gut lymph nodes, in part reflecting the challenges of imaging these anatomically intricate structures during early life. Intriguingly, neonatal mesenteric lymph nodes possess enhanced capacity to support pTreg differentiation that depends on stromal rather than immune cells^39^, suggesting that the developing lymph node microenvironment actively contributes to the establishment of tolerance. However, how this tolerogenic microenvironment is spatially organized, where functionally distinct cDC and TC subsets are positioned, and how they gain access to dietary and microbiota-derived antigens during early life remain unknown.

Here, we generated complementary single-cell-resolution Xenium and whole-transcriptome Visium HD spatial atlases of small intestine- and colon-draining lymph nodes during weaning and adulthood. We define developmentally distinct lymph node architecture, cytokine environments and specialized APC niches, including an early life TC I-enriched subcapsular compartment lacking classical antigen-relay macrophages coincident with suppressed follicular responses, and a TC IV niche associated with RORγt⁺ pTreg cell priming. We further identify specialized intestinal TCs and establish tissue-to-node migration by TC III and IV. Together, this atlas provides a spatial framework for understanding how lymph node architecture, antigen access and APC specialization shape intestinal tolerance and immunity across developmental stages and intestinal regions.

## RESULTS

### A single-cell spatial atlas of gut-draining lymph nodes during weaning and adulthood

The peri-weaning period (in mice postnatal day (P)12-18) represents a critical transition window in which introduction of solid food not only leads to exposure to a diverse array of dietary antigens but also an increase in the abundance and diversity of the gut microbiota. Food- and microbiota-specific pTreg cells appear in gut lymph nodes at ∼P14 and accumulate rapidly thereafter^8,15^. Thus, to establish the cellular architecture and microenvironment of gut-draining lymph nodes during the establishment of tolerance to intestinal antigens and their developmental-stage specific features, we collected mesenteric lymph node (mLN) chains from peri-weaning (P14) and adult (6-week-old) C57Bl/6 mice (n = 3 mice per age group) (Figure 1A). Samples encompassed small intestinal lymph nodes (siLNs) draining the proximal-to-distal small intestine (duodenum, jejunum, ileum) and colonic lymph nodes (cLNs). We profiled adjacent tissue sections using complementary spatial transcriptomic platforms: a custom 472-gene 10x Genomics Xenium panel designed to finely resolve immune and stromal cell identities, cell states and selected ligand–receptor pairs, and Visium HD for broader transcriptome-wide coverage. Two adjacent sections were analyzed by Xenium, with a third adjacent section profiled by Visium HD.

**Figure 1.**
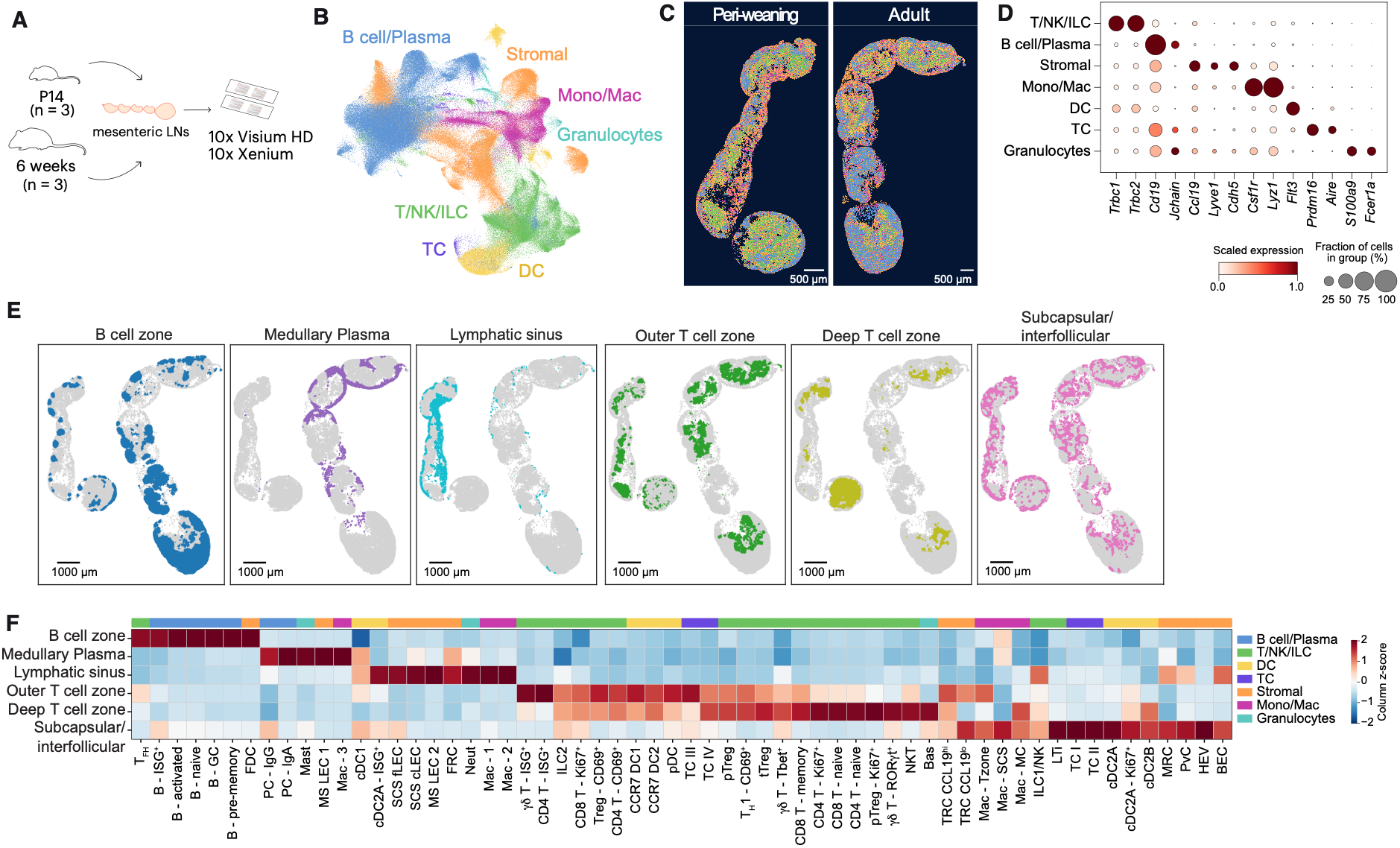
Single-cell spatial atlas resolves the cellular and anatomical niches of peri-weaning and adult gut-draining lymph nodes. (A) Schematic showing experimental design. Mesenteric lymph node (mLN) chains were collected from postnatal day (P)14 and 6-week-old mice (n = 3 per group), sectioned, and profiled with 10x Genomics Xenium (472-gene custom panel) and Visium HD. (B) Uniform manifold approximation and projection (UMAP) of cells within the Xenium dataset, colored by broad cell lineage: B cell/Plasma, T/Natural Killer (NK)/innate lymphoid cell (ILC), monocytes/macrophages (Mono/Mac), stromal cells, dendritic cells (DC), Thetis cells (TC), and granulocytes. (C) Spatial overlays of representative mLN chains from P14 (peri-weaning) and 6-week-old (adult) mice. Each cell is colored by broad cell lineage annotation. Scale bars, 500 µm. (D) Dot plot with relative expression of selected marker genes for each broad lineage. (E) Representative peri-weaning and adult mLN, colored by niche. (F) Heatmap of the relative abundance of fine cell types (columns) in each niche (rows).

Cell segmentation and transcript assignment of Xenium data using segger^40^ generated spatially resolved single-cell expression profiles for 1,267,407 cells across all samples. Unsupervised clustering identified the major lymph node compartments, including stromal and vascular cells, monocytes and macrophages, DCs, TCs, T, natural killer (NK) and innate lymphoid cells (ILCs), B and plasma cells, and granulocytes (Figures 1B –1D). Cell-type composition and gene detection were highly reproducible across adjacent tissue sections, with similar cellular profiles observed across three independent lymph node chains at each age (Figures S1A–S1C). The siLN portion of adult replicate 3 exhibited reduced cell density and was therefore excluded from downstream spatial analyses.

Subclustering within major immune and stromal lineages resolved a rich diversity of immune and stromal populations (Figures S2A–S2G). Among lymphoid cells, we identified RORγt^+^Foxp3^+^ (pTreg) and RORγt^−^Helios(*Ikzf2*)^+^Foxp3^+^ thymic (t)Treg cells, T_FH_ and T_H_1 cells, activated and proliferating T cell states, innate lymphoid cells (ILC) (Natural Killer (NK)/ILC1, ILC2), LTi cells, RORγt and T-bet^+^ γ8 T cells, and NKT cell subsets (Figure S2A). B cells encompassed naive, activated, memory and germinal center (GC) B cells, together with IgA^+^ and IgG^+^ plasma cells (Figure S2B). Stromal subclustering resolved high endothelial venule (HEV) and other blood endothelial cells (BECs), CXCL12^+^ fibroblastic reticular cells (FRCs), lymphoid tissue organizer-derived RANKL^+^ marginal reticular cells (MRCs), follicular dendritic cells (FDCs), CCL19^hi^ and CCL19^lo^ T-zone reticular cells (TRCs), and four lymphatic endothelial cell (LEC) populations encompassing subcapsular sinus ceiling and floor LECs and medullary sinus LECs (Figure S2C). APCs included cDC1, cDC2A and cDC2B, their CCR7^+^ activated or migratory counterparts, plasmacytoid DCs (pDCs) (Figure S2D), and TC I–IV (Figure S2E). In addition, we resolved anatomically distinct macrophage populations (Figure S2F) alongside granulocyte populations comprising neutrophils, basophils and mast cells (Figure S2G).

We next characterized multicellular organization, using spatial community analysis to identify recurrent niches with characteristic cell-type composition. This revealed 6 major niches corresponding to distinct anatomical regions (Figures 1E–1F): a medullary plasma cell niche enriched for medullary sinus LECs, macrophages and plasma cells; B cell follicles comprising naïve, activated, memory and GC B cells, T_FH_ cells and FDCs; a lymphatic sinus niche comprising subcapsular sinus and medullary LECs, neutrophils and FRCs; the interfollicular and subcapsular region enriched for MRCs, LTi cells and SCS macrophages; the outer T-cell zone comprising CCR7^+^ cDC1, CCR7^+^ cDC2 and CCL19^+^ FRCs, and finally the deeper T-cell zone, enriched for CD4 and CD8 T cell subsets (Figure 1E for a representative lymph node and Figure S3A for all lymph nodes).

Analysis of receptor-ligand pairs captured by our Xenium panel revealed spatially enriched interactions across distinct cellular niches, including chemokine signaling pathways that shape lymph node architecture (Figure S3B). Notably, CCL11-CCR3 was enriched in medullary plasma cell niches, with FRCs representing the predominant source of *Ccl11* and IgA plasma cells expressing the highest levels of *Ccr3* (Figure S3C), findings independently supported by ImmGen bulk RNA-seq datasets (Figure S3D). Previous studies have identified specialized FRCs within the medulla (medRCs) that support plasma cell migration and survival^41^. Our spatial analysis revealed close associations between *Ccl11^+^* FRCs and *Ccr3^+^* IgA plasma cells within the medulla (Figures S3E and S3F), identifying a previously unrecognized chemokine axis that may contribute to plasma cell positioning or retention. By contrast, CXCL12–CXCR4 interactions were enriched in subcapsular/interfollicular regions, highlighting distinct chemokine environments that support gut lymph node architecture.

Together, these analyses establish a spatially resolved cellular and molecular reference atlas for gut-draining lymph nodes, defining their major anatomical niches and identifying candidate stromal–immune interactions that may regulate their organization and function across developmental stages and intestinal drainage regions.

### Developmental and regional specialization of gut-draining lymph nodes

To determine whether the lymph node architecture differs between peri-weaning and adult mice, we compared niche abundance as well as the cellular composition within each defined niche. Adjacent sections of the same biological lymph node chain displayed reproducible niche composition (Figure S4A), and were thus pooled for comparisons across biological replicates. Notably, peri-weaning mLN chain 2 was dominated by medullary sinus niches, likely reflecting the plane and depth of sectioning, which preferentially captured the medullary compartment. Adult lymph nodes were enriched for B cell, plasma cell and outer T cell zone niches, whereas peri-weaning lymph nodes were dominated by the lymphatic sinus, subcapsular sinus, and T cell zone niches (Figure 2A), suggesting prominent trafficking of antigen and T cell surveillance during early life. Developmental differences extended beyond niche abundance, with anatomically equivalent niches exhibiting distinct cellular compositions in peri-weaning and adult lymph nodes (Figure 2B). In peri-weaning mice, the outer T cell zone niche was enriched for tolerogenic TC IV, ILC2s, NK and γ8 T cells, whereas the corresponding adult niche was enriched for T-cell zone macrophages. Developmental differences were also evident in the subcapsular sinus and interfollicular niches, with enrichment of TC I, TC II and LTi cells during peri-weaning compared with cDC2A and cDC2B subsets in adulthood.

**Figure 2.**
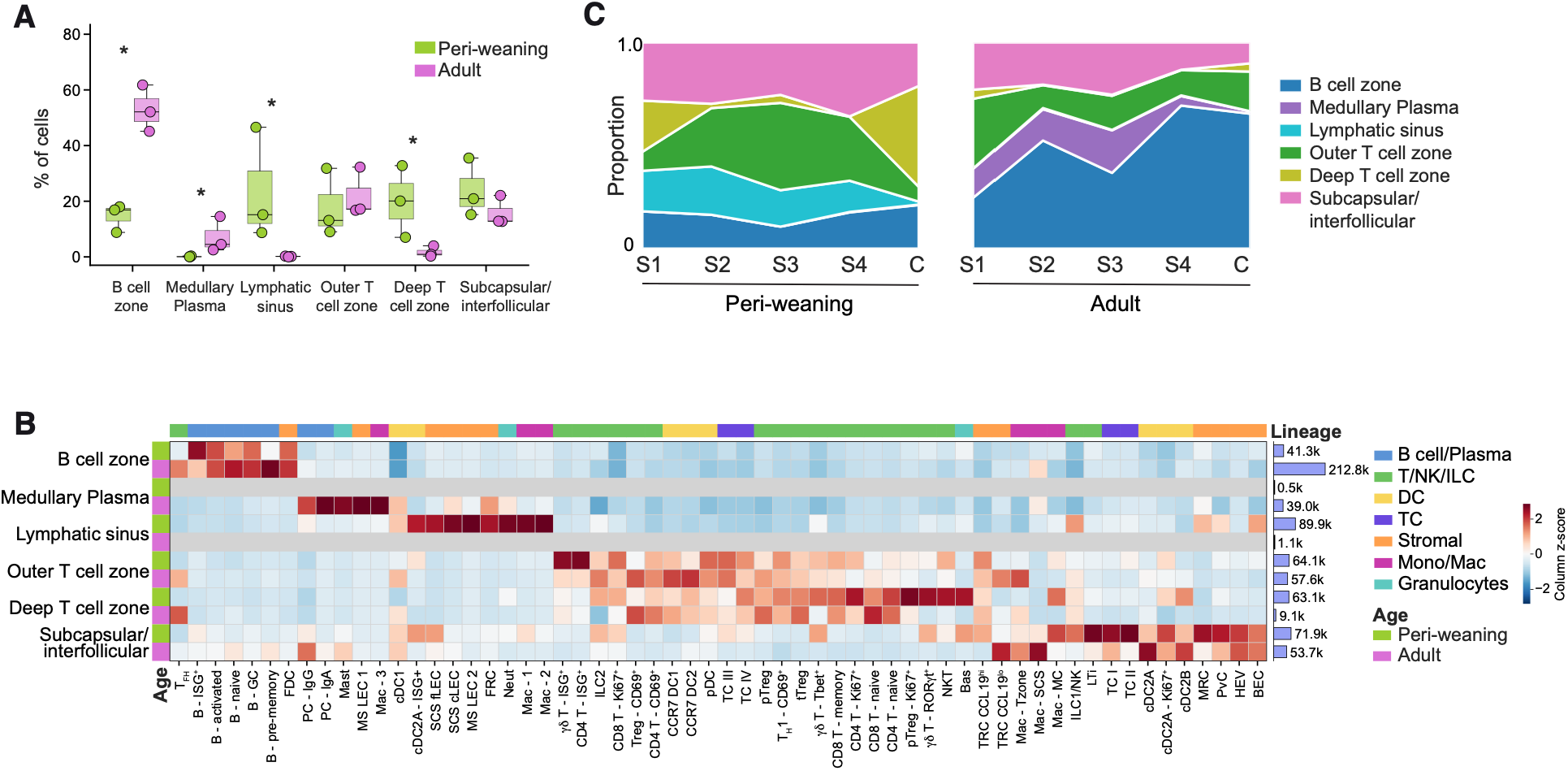
Developmental stage and intestinal drainage region shape lymph node architecture. (A) Percentage of cells assigned to each spatial niche in peri-weaning (P14) and adult (6-week-old) mice (n = 3 per age group). (B) Heat map of relative cell type abundance within each niche in peri-weaning or adult mice. (C) Niche proportions along the lymph node chain in peri-weaning (left) and adult (right) mice. Nodes are ordered from proximal to distal (small intestinal S1-S4, colonic (C)). Box plots in A show the median and interquartile range (IQR); limma t-test. *FDR < 0.05.

Previous studies have demonstrated anatomically segregated lymphatic drainage from the small and large intestine, accompanied by regional differences in immune composition^42^. Individual lymph nodes within the mesenteric chain of adult mice could be readily distinguished. However, at P14, the chain comprises a discrete colon-draining node and a more continuous region of small intestine-draining lymphoid tissue. We therefore partitioned the siLN chain into four segments (S1–S4) ordered from proximal to distal intestinal drainage, guided by anatomical features (Figure S4B), and analyzed these distinct regions. Two areas were excluded from regional analyses: a P14 chain with discontinuous siLN tissue and a partially captured adult cLN. Principal component analysis revealed broadly concordant profiles among corresponding regions across biological replicates, supporting the reproducibility of our regional assignments (Figure S4C).

Within each age group, niche composition analysis revealed that proximal to distal siLN regions were similar to each other but distinct from their cLN counterparts (Figure 2C). The most striking difference between cLNs and siLNs was the marked reduction in the medullary plasma niche in adult cLNs and the corresponding reduction in the medullary lymphatic sinus niche in peri-weaning cLNs (Figure 2C). Analysis of individual cell types confirmed that this reflected a global paucity of all medullary sinus-associated cell types within cLN including medullary LECs, macrophages and plasma cells (Figure S4D). This finding is in line with recently reported reduced expression of medullary sinus stromal and macrophage associated transcripts in bulk RNA-seq on whole human and mouse cLNs relative to siLNs^43^, indicating that region-specific differences in our spatial dataset were not due to profiling of a non-representative section of the lymph node. The distinct sinus-associated architectures of siLNs and cLNs, suggests different capacities for handling lymph-borne antigens, with potentially important consequences for immune responses to intestinal antigens.

To further define regional differences in gut lymph node function, we performed differential gene expression analysis of siLNs and cLNs from peri-weaning and adult mice. Peri-weaning cLNs were enriched for Treg-associated genes (*Foxp3, Ctla4, Il2ra, Stat5a*) whereas adult cLNs exhibited greater expression of GC-associated transcripts (*Bcl6, Cr2, Gcsam*) (Figure S4E). Peri-weaning siLNs were enriched for lymphatic endothelial (*Prox1, Lyve1*) and sinus-associated macrophage (*Fcgr1, C5ar1*) genes, whereas adult siLNs displayed increased expression of IgA plasma cell-associated genes (*Igha, Jchain*), together with *Aldh1a2*, encoding a key enzyme in retinoic acid synthesis. Given the established role of retinoic acid in promoting IgA class switching in B cells, the developmental enrichment of *Aldh1a2* may contribute to the shift from early-life commensal-specific IgG responses towards IgA-dominated responses after weaning^44,45^.

Together, these findings identify developmental stage and intestinal drainage region as major determinants of gut lymph node organization. Persistent regional differences in sinus architecture are accompanied by pronounced developmental changes in cellular composition and immune activation, suggesting that anatomically distinct gut lymph nodes undergo specialized programs of immune maturation.

### Peri-weaning interferon responses in small intestinal lymph nodes

To investigate the spatially coordinated transcriptional programs underlying developmental differences in gut lymph nodes, we leveraged the broader transcriptome captured by Visium HD (∼19,000 genes) and applied spatial Hotspot to identify genes with correlated expression^46^. This analysis resolved 24 spatial gene modules corresponding to cellular niches, immune activation and cytokine signaling (Figures 3A and S5A–S5B). Comparison of module scores across ages revealed striking differences in the cellular and cytokine environments of peri-weaning and adult lymph nodes (Figure S5C). Peri-weaning nodes were enriched for a macrophage–stromal module (M1; *Marco, Siglec1, Csf1r, C1qa, Lyve1, Dcn, Cldn5*), and an HEV-associated lymphocyte homing program (M6; *Glycam1, Ubd, Madcam1, Ackr1*) reflecting a dense lymphatic-vascular network for antigen trafficking and T cell surveillance. Notably, two interferon-associated modules were enriched within the T cell zones of peri-weaning lymph nodes: M7, comprising interferon-stimulated genes (ISGs) and CCR7^+^ cDC1- and cDC2-associated genes (*Flt3, Il12b, Cd1d1*), and M9 containing ISGs, T cell activation genes (*Lcp2, Itm2a, Tnfrsf1b*) and CCR7^+^ cDC-associated genes (*Marcksl1, Cxcl16*), suggesting spatially coordinated IFN sensing, cDC activation and T cell priming within peri-weaning gut lymph nodes.

**Figure 3.**
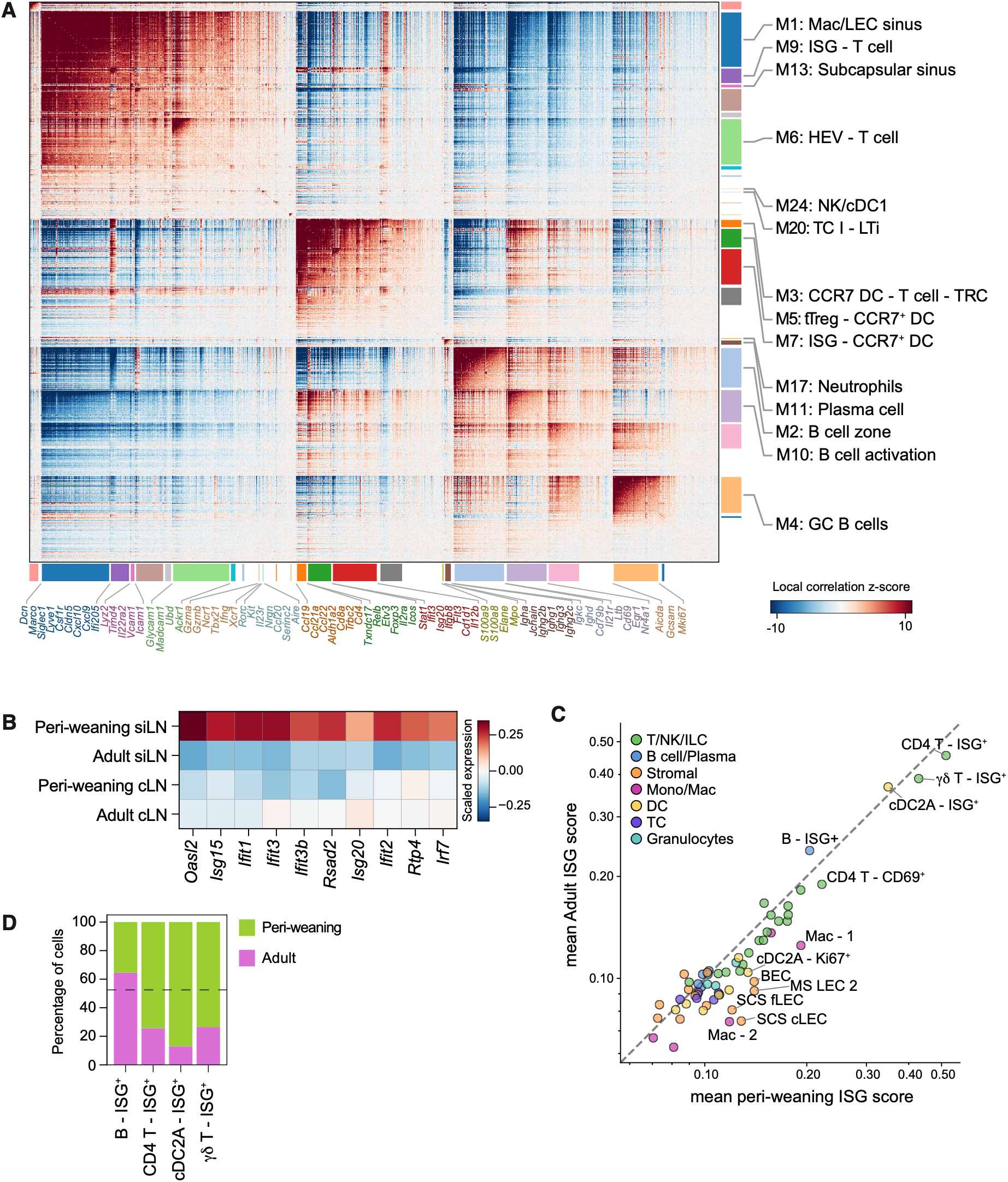
Peri-weaning lymph nodes exhibit spatially coordinated interferon-responsive programs. (A) Hotspot analysis of the 19,045 genes detected in Visium HD data. The spatial correlation matrix of gene pairs and hierarchical clustering define 24 modules with ≥30 genes per module. (B) Heat map of differentially expressed interferon-stimulated genes (ISG) across peri-weaning and adult siLNs and cLNs. (C) Relative expression of ISGs (as in B) for each fine cell type in adult (y) or peri-weaning (x) mice. (D) Proportion of adult and peri-weaning cells in each ISG⁺ cluster. Dashed line indicates the overall proportion of adult and peri-weaning cells across all cells.

Analysis of the Xenium dataset confirmed enrichment of ISGs in the siLNs of peri-weaning mice (Figure 3B). Across multiple immune and stromal lineages, ISG expression was elevated in peri-weaning cells relative to their adult counterparts, most notably within macrophages and LECs associated with lymphatic sinuses (Figure 3C). This spatial distribution raises the possibility that interferons or upstream IFN-inducing stimuli reach gut lymph nodes through intestinal lymphatics. In addition to the increase in ISG expression on a per cell basis, the frequency of B cells, CD4 T cells and cDC2s displaying high levels of ISGs was increased in peri-weaning lymph nodes (Figure 3D). Thus, an interferon-associated response is a widespread property of the developing gut lymph node rather than a program confined to one particular cell type. In keeping with the spatial enrichment of ISGs within the T cell zone, peri-weaning lymphoid cells expressed higher levels of both *Ifnar1* and *Ifngr1* (Figure S5D), suggesting the potential for responsiveness to type I (IFN-α/β) and type II (IFN-γ) interferons, whose downstream transcriptional programs substantially overlap. Notably, this response was already evident at P14, preceding the previously described IFN-γ-associated “weaning reaction” in the small intestinal lamina propria of P21 mice that coincides with physical weaning of mice from their mothers^47^.

In contrast to the pronounced interferon-responsive program in peri-weaning nodes, analysis of differentially expressed cytokine signaling genes revealed enrichment of *Il21*, reflecting mature T_FH_ responses, as well as a modest increase in type 3-associated immune-response genes in adult lymph nodes, including *Il6* and *Il17* (Figure S5E). Among the most highly upregulated genes in adult lymph nodes was *Il22ra2*, encoding the soluble IL-22 binding protein (IL-22BP), a potent antagonist of IL-22. Analysis of the Xenium cell atlas identified subcapsular sinus macrophages (SSMs) as the sole source of *Il22ra2*, with expression restricted to adult nodes (Figure S5F). Although IL-22BP expression by lymph node SSMs has not previously been described, analysis of the GutPath mesenteric lymph node atlas confirmed this pattern^48^, revealing a previously unrecognized feature of SSMs. By contrast, LTi cells, localized adjacent to SSMs, were the dominant source of IL-22 at both ages (Figure S5G) and were far more abundant in peri-weaning lymph nodes (Figure S5H). Thus, peri-weaning nodes combine an expanded population of LTi cells expressing Il22 with an absence of detectable Il22ra2 expression, whereas adult nodes exhibit the inverse configuration, with fewer Il22-expressing cells and increased SSM-derived Il22ra2. These findings reveal reciprocal developmental regulation of IL-22 and its antagonist IL-22BP, suggesting that the balance between IL-22 production and antagonism shifts substantially during postnatal lymph node maturation. At present the role of lymph node LTi-derived IL-22 is not known. Besides the established role of IL-22BP in regulating epithelial antigen uptake in Peyer’s patches^49^, IL-22 has been implicated in stromal CXCL13 production during tertiary lymphoid tissue formation^50^. Whether the developmentally regulated IL-22–IL-22BP axis influences lymph node organization or instead acts at distal intestinal epithelial sites remains to be determined.

Together, these findings reveal a profound developmental reorganization of the gut lymph node cytokine environment, establishing distinct immunological contexts for the active priming and differentiation of T cells during early life and the maintenance of immune homeostasis in adulthood.

### A TC I-enriched early-life subcapsular niche lacks classical antigen-relay macrophages

Intestinal bacteria and bacterial products have been reported to reach mesenteric lymph nodes during early postnatal life^51,52^; however, the cells that capture these antigens and the pathways through which they are distributed have not been defined. The subcapsular sinus represents a major entry point for small soluble lymph-borne proteins into the lymph node conduit network and medullary sinuses. Larger soluble antigens, immune complexes and microbial particulate antigens are captured by specialized subcapsular macrophages, which facilitate transfer to FDCs for subsequent presentation to follicular B cells, thus supporting sustained humoral responses^32,33^. While age-dependent differences in macrophages and capture of bacterial antigens have been reported in the spleen^53^, the timing of antigen-acquisition programs in early life gut lymph nodes has not been established.

We first examined the identity, abundance and spatial distribution of sinus macrophages, given their key role in antigen uptake and transfer. Integration of published gut lymph node macrophage transcriptional signatures with spatial localization resolved classical subcapsular sinus macrophages (SSMs; *Siglec1^+^Timd4^+^*), medullary cord macrophages (MCMs; *Adgre4^+^Ace^+^*), T-cell zone macrophages and three *Marco^+^Siglec1^+^* macrophage states (Mac 1–3) (Figures 4A–4C and S1F)^43^. Postnatal SSMs and medullary sinus macrophages (MSMs) differentiate from bone-marrow derived monocytes that enter the lymph node^54^. Pseudotime analysis ordered Mac 1–3 cells along a developmental continuum, with progressive acquisition of the canonical MSM program (Figures 4D and 4E), suggesting that Mac 1 represented the earliest monocyte precursors. Notably, Mac 1 and Mac 2 cells were almost exclusively derived from peri-weaning nodes (Figures 4C, 4D and 4F). Mac 2 expressed proliferation-associated genes, consistent with an early stage of MSM differentiation, but exhibited reduced expression of the antigen-binding receptors *Siglec1* (CD169) and *Marco*, and lacked *Cd209b* (SIGN-R1), a key receptor for bacterial polysaccharide capture^55^ (Figures 4E and 4G). Mac 3 cells were predominantly found in adult lymph nodes where they localized to the medullary sinus niche, confirming their MSM identity (Figures 4C and 2B). Unexpectedly, Mac 1 and 2, despite exhibiting an MSM-like phenotype, occupied not only medullary sinuses but also subcapsular regions of peri-weaning nodes, directly adjacent to floor scLECs, where macrophages bearing the conventional “adult” SSM phenotype were markedly depleted (Figures 4C and 4H). Thus, the peri-weaning subcapsular sinus is characterized by a distinct population of MSM-like macrophages, albeit with reduced or absent expression of key antigen-binding receptors, replacing the classical SSM population found in adults. Given the established roles of SSMs in capturing and relaying intact antigens to follicular B cells and FDCs, and the role of CD209b^+^ MSMs in the recognition of microbial polysaccharides^32,33,55^, these findings suggest developmental differences in the capacity of sinus macrophages to capture and distribute lymph-borne antigens.

**Figure 4.**
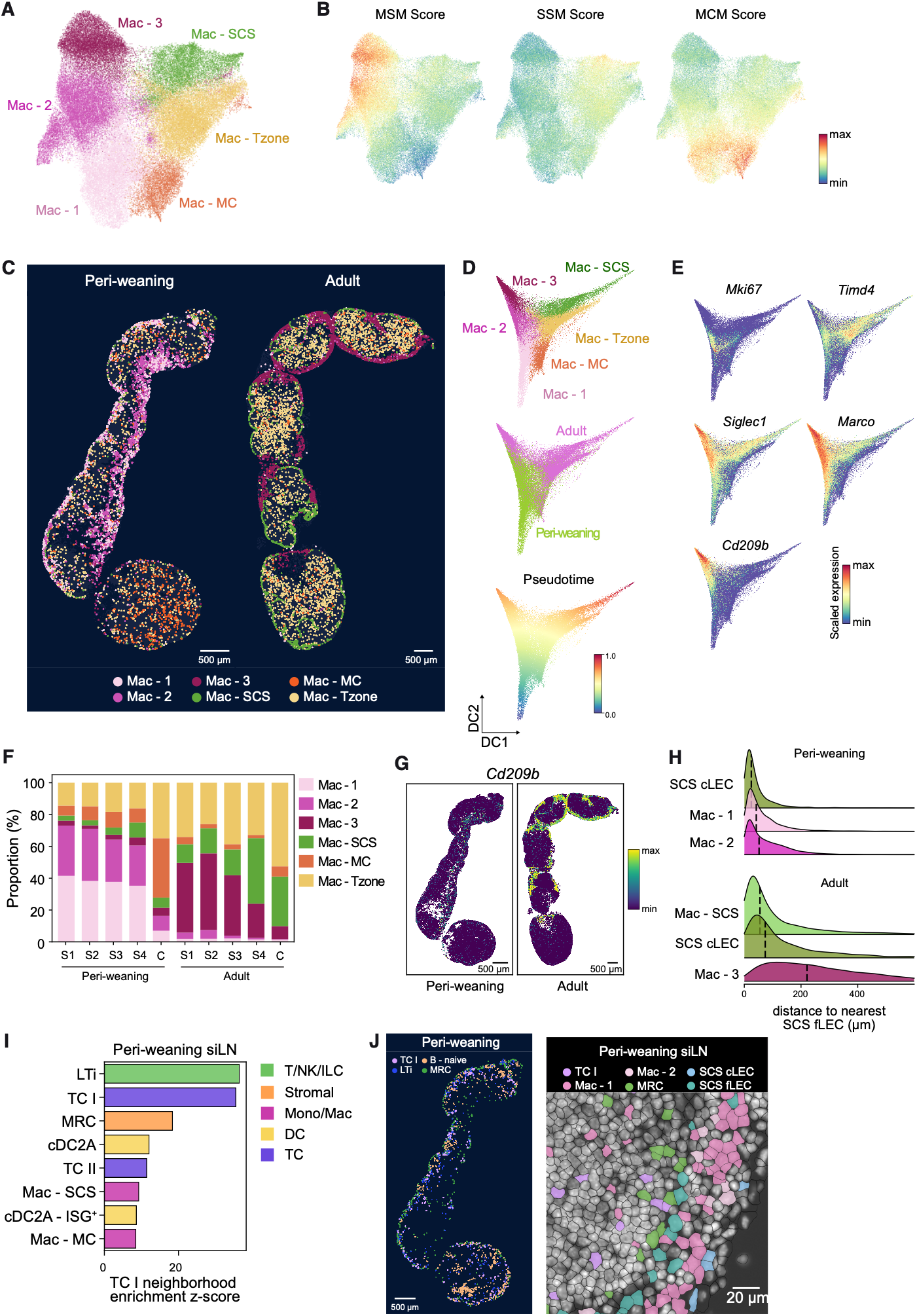
A TC I-enriched early-life subcapsular niche lacks classical antigen-relaying macrophages. (A) UMAP of macrophages colored by fine cell type annotation. (B) UMAP colored by gene signature scores for medullary sinus (MSM), subcapsular sinus (SSM), and medullary cord (MCM) macrophages, derived from published bulk RNA-seq on gut lymph node macrophages (Fatkhullina et al.). (C) Spatial overlays of macrophage subsets in representative peri-weaning and adult lymph node chains, colored by fine cell type annotation as in (A). Scale bars, 500 µm. (D) Diffusion map embedding of macrophages colored by cell type (top), age (middle), or Palantir pseudotime (bottom). (E) Diffusion map embedding colored by expression of selected genes. (F) Proportion of each macrophage subset among total macrophage cells across siLNs (S1-S4, proximal to distal) and cLNs of peri-weaning and adult mice. (G) Spatial expression of *Cd209b* in representative peri-weaning and adult mLNs. Scale bars, 500 µm. (H) Distribution plots of the distance from each cell in the indicated source populations to the nearest subcapsular sinus floor lymphatic endothelial cell (SCS fLEC) in peri-weaning (top) and adult (bottom) siLNs. Dashed lines indicate median values. (I) Neighborhood enrichment of cell types around TC I in peri-weaning siLN. (J) Spatial distribution of TC I, lymphoid tissue inducer (LTi), marginal reticular cells (MRC) and naive B cells in a representative peri-weaning mLN chain (left; scale bar, 500 µm) and a magnified region of the siLN (right) showing indicated cell types. Scale bar, 20 µm.

The differentiation and maintenance of SSMs depend on spatially coordinated signals within the subcapsular niche, including CSF1 from subcapsular sinus LECs, RANKL from MRCs, and lymphotoxin (LTα1β2) from naïve B cells, whose expression is promoted by FDC-derived CXCL13^33,54,56–58^. The MSM-like phenotype of peri-weaning subcapsular macrophages was reminiscent of the SSM-to-MSM phenotypic diversion observed following reduced RANK or LTβR signaling^57^, prompting us to examine these pathways. Peri-weaning nodes exhibited a diminished CXCL13–CXCR5 FDC–B cell axis (Figure S6A and S6B), potentially limiting B cell-derived lymphotoxin. While RANKL^+^ MRCs are enriched in peri-weaning nodes (Figures 2B and S2B), receptor-ligand analysis identified TC I as a potential source of osteoprotegerin (OPG), a decoy receptor for RANKL that may locally attenuate RANK signaling (Figures S6A–S6C). Consistent with this, TC I were enriched in the subcapsular sinus, in direct contact with MRCs and sinus macrophages (Figures 2B and 4I–J).

Together, these findings define an early-life subcapsular niche characterized by TC I enrichment and predominance of MSM-like macrophages in place of classical SSMs, potentially reflecting altered RANK and/or LTβR signaling. Given the established role of classical SSMs in antigen capture and relay, this developmental organization may limit the delivery of intact antigens to the follicular compartment, thereby contributing to attenuated T_FH_ maturation and T-dependent B cell responses during the establishment of intestinal tolerance.

### Attenuated T_FH_ and enhanced pT_reg_ differentiation within peri-weaning colonic lymph nodes

Consistent with this possibility, T_FH_ cells and activated, memory and GC B cell populations were relatively scarce at P14 compared with adult nodes (Figures 5A–D). However, diminished T_FH_ responses did not reflect a global reduction in T cell activation. Rather, peri-weaning lymph nodes displayed a striking enrichment of recently activated (CD69^+^) and proliferating (Ki67^+^) CD4 and CD8 T cells within the T-cell zone (Figures 5A and 5F), consistent with extensive T cell activation and proliferation during this window of increased exposure to intestinal antigens. Most notably, peri-weaning cLNs were highly enriched for Ki67^+^ pTreg cells (Figure 5D), consistent with the developmental wave of food- and microbiota-specific pTreg cell differentiation during this period. Whereas tTreg cells were equally abundant within the APC-rich outer T cell zone and deep T cell zone, Ki67^+^ RORγt⁺ pTreg cells were almost exclusively present in the deep T cell zone (Figure 2B), suggesting differentiation within a distinct niche. Besides Treg cells, peri-weaning colonic nodes were also enriched for recently primed T_H_1 cells (IL-2^+^CD69^+^IFN-γ^+^), albeit T_H_1 cells were proportionally less abundant than pTreg cells (Figure 5E).

**Figure 5.**
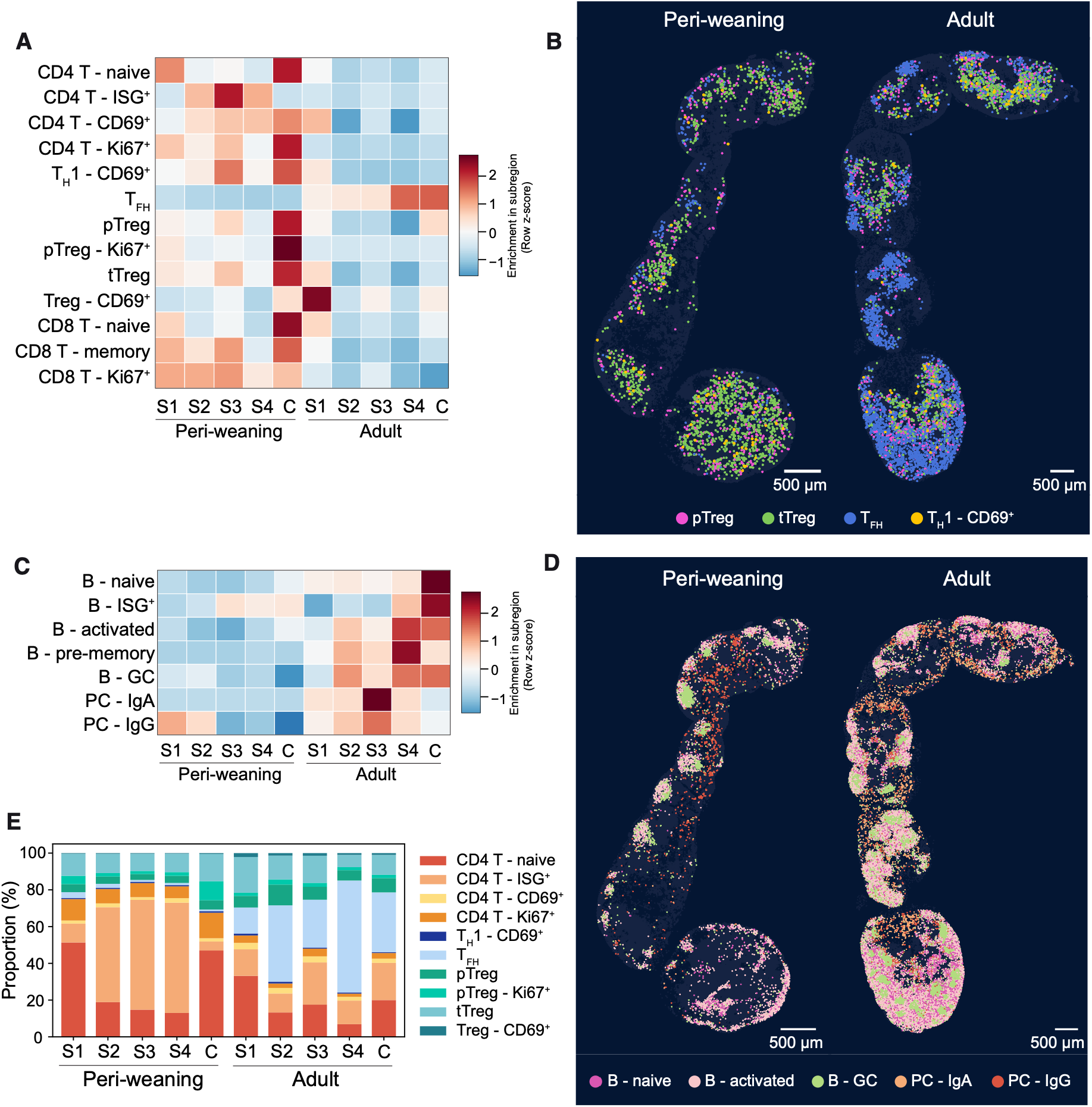
Peri-weaning colonic lymph nodes support pTreg cell differentiation. (A) Heatmap of the relative abundance of CD4^+^ and CD8^+^ T cell subsets (rows) in each mLN subregion. (B) Spatial overlays of indicated T cell types in representative peri-weaning and adult mLNs. (C) Heatmap of the relative abundance of B cell and plasma cell (PC) subsets in each mLN subregion (germinal center B cell; B – GC). (D) Spatial overlays of indicated B cell types in representative peri-weaning and adult mLNs. (E) Proportion of CD4^+^ T cell subsets among total CD4^+^ cells in each subregion across ages.

Beyond the reduced abundance of GC and memory B cells, peri-weaning lymph nodes exhibited poorly developed B cell follicles with markedly fewer naïve B cells (Figures 5C and 5D). Nevertheless, plasma cells were distributed throughout the medullary compartment and were predominantly IgG⁺, in contrast to the IgA-dominated plasma cell population in adult lymph nodes (Figures 5C and 5D). This developmental shift in plasma cell isotype composition is consistent with previous studies demonstrating that commensal-specific IgG responses are established during early life^44,59^.

Together, these findings reveal that early-life gut lymph nodes are not simply immature or quiescent, but sites of extensive T cell activation and proliferation, dominated by pTreg cell differentiation. Adult nodes, by contrast, exhibited more developed T_FH_, GC B cell and plasma cell-associated programs, alongside fewer T cells in recently activated or proliferative states. This developmental shift raises the possibility that restricted follicular antigen access and B cells, together with the specialized APC niches of peri-weaning nodes, favors the establishment of intestinal tolerance while limiting T_FH_-dependent humoral responses which are known to suppress pTreg cells^88^.

### Spatial compartmentalization of specialized APC subsets during early-life tolerance induction

The enrichment of both recently primed RORγt⁺ pTreg cells and T_H_1 cells in peri-weaning colonic lymph nodes raised the question of whether distinct APC niches within the T cell zone of peri-weaning lymph nodes preferentially support distinct CD4 T cell fates. Given the established roles of cDC and TC subsets in directing effector vs regulatory CD4 T cell differentiation, we first examined their relative abundance and spatial organization. In contrast to cDC subsets, which exhibited no uniform age-dependent changes in abundance, all TC subsets were enriched in peri-weaning siLNs and cLNs (Figures 6A and 6B), a finding confirmed by flow cytometric analysis of TCs in individual siLNs and cLNs of P14 and adult mice (Figure 6C). Although the developmental enrichment of pTreg-inducing TC IV may contribute to the enhanced Treg responses during early life, these cells remained considerably less abundant than CCR7⁺ cDC1 and cDC2. We therefore asked whether their spatial positioning within specialized cellular niches might further contribute to their distinct functions during early-life tolerance induction.

**Figure 6.**
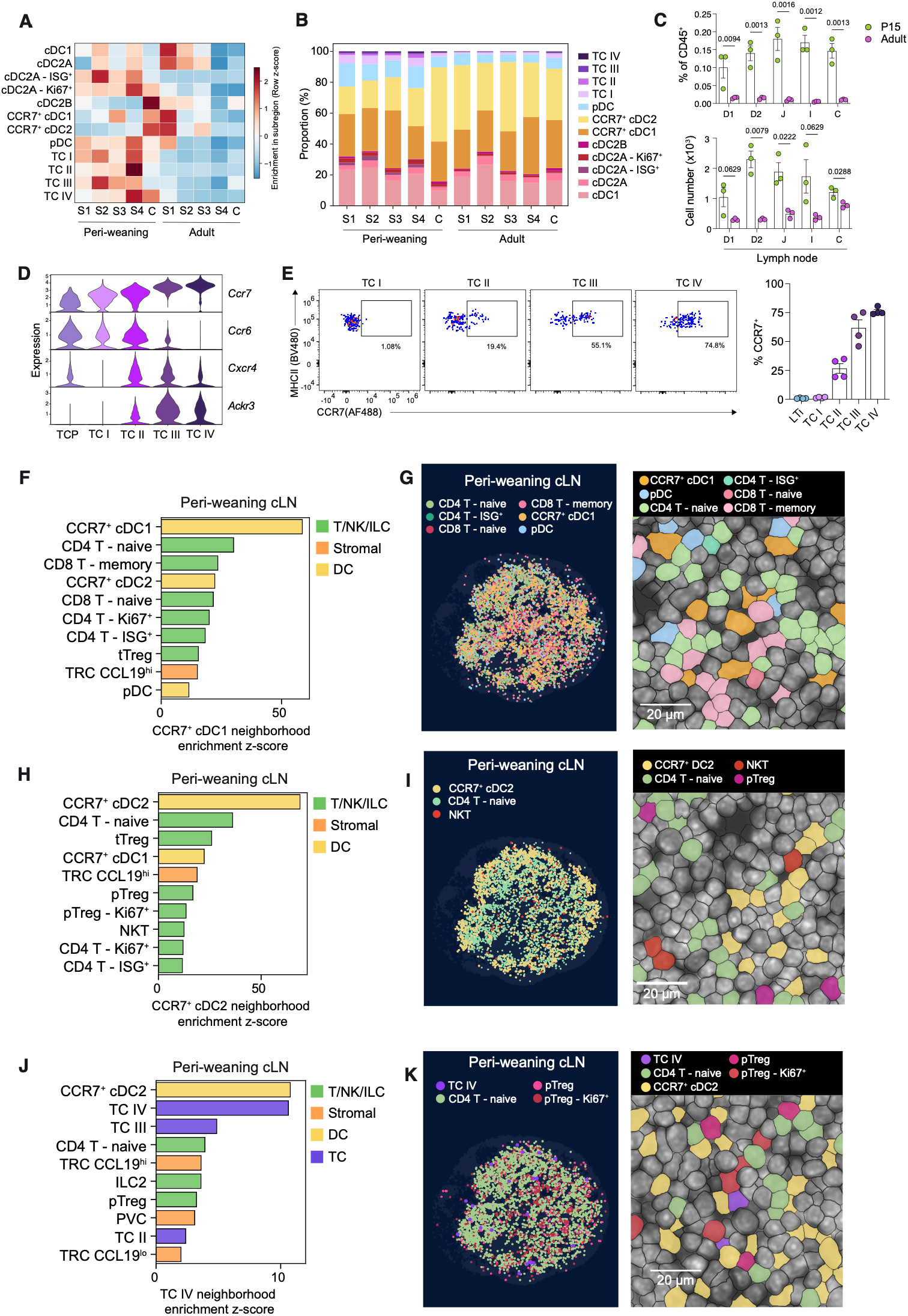
Specialized dendritic cell and Thetis cell subsets occupy distinct APC–T cell niches. (A) Heat map of the relative abundance of DC and TC subsets (rows) in each lymph node subregion (columns) of peri-weaning and adult mice, ordered from proximal to distal. (B) Proportion of DC and TC subsets among DCs and TCs in each subregion, shown as stacked bar plots. (C) Percentage and number of TCs in individual lymph nodes of mLNs of P15 and 13-week *Rorc^Venus-creERT^*^2^ mice (n = 3). Gut lymph nodes: D1, portal; D2, distal duodenum; J, jejunum; I, ileum. (D) Expression of selected chemokine receptors by TCs profiled by scRNA-seq (Cabric et al.). (E) Representative flow cytometry of CCR7 expression by TC subsets in mLNs of P17 *Rorc^Venus-creERT^*^2^ mice and summary graph (n = 4). Data in (C) and (E) are representative of three independent experiments. Error bars: mean ± s.e.m.; multiple unpaired t-tests (C). All *P* values are indicated on the corresponding graphs. (F, H, J) Neighborhood enrichment in peri-weaning cLNs around CCR7⁺ cDC1 (F), CCR7⁺ cDC2 (H), and TC IV (J). Bars show the neighborhood enrichment z-score for the cell types with the highest enrichment, colored by broad lineage. (G, I, K) Spatial overlays of the cell types enriched around CCR7⁺ cDC1 (G), CCR7⁺ cDC2 (I), and TC IV (K) in a representative peri-weaning cLN (left) and magnified regions (right).

Mapping APC subsets onto the cellular neighborhoods defined above revealed pronounced spatial segregation. Resident cDC1s were broadly distributed across subcapsular/interfollicular, medullary and outer T cell zone niches, whereas cDC2A and cDC2B were concentrated in subcapsular and interfollicular regions (Figures 1F and 2B). By contrast, CCR7⁺ DC1 and DC2 accumulated in the CCL19⁺ reticular cell-rich outer T cell zone (Figures 1F and 2B). Within the T cell zone, CD4 and CD8 T cells occupied distinct but overlapping territories, with CD8 T cells concentrated in the deeper paracortex and CD4 T cells more abundant in peripheral regions (Figure S7A). Notably, CCR7⁺ DC1 were distributed across both CD4- and CD8-enriched regions, whereas CCR7⁺ DC2 were predominantly restricted to the CD4-rich outer T cell zone (Figure S7B). This organization is consistent with the established roles of CCL19^hi^ TRCs in maintaining central CD8 T cell niches, and GPR183-dependent oxysterol gradients in positioning CD4 T cells and CCR7^+^ cDC2s towards the outer T cell zo^60,61^. Notably, peri-weaning CCR7^+^ cDC1s expressed higher levels of IL-12 (Figure S7C), consistent with increased CCR7^+^ cDC activation and licensing by interacting T cells and type I cytokines within peri-weaning lymph nodes, as suggested by our earlier hotspot analysis.

TC subsets were similarly compartmentalized: TC I and II preferentially occupied subcapsular and interfollicular regions, whereas TC III localized to the outer T cell zone, and TC IV was equally abundant in the inner and outer T cell zone (Figure 2B). These distributions were consistent with distinct chemokine receptor programs: TC III and IV expressed CCR7 transcript and protein, whereas TC I lacked detectable expression and TC II expressed low levels (Figures 6D and 6E). Conversely, TC I and II express CCR6 (Figure 6D), whose ligand CCL20 was enriched in subcapsular sinus floor LECs (Figures S2C and S7D). TCs also expressed *Ackr3* (Figure 6D), encoding the atypical chemokine receptor ACKR3/CXCR7, which scavenges extracellular CXCL12 to maintain chemokine gradients and preserve CXCR4 responsiveness.

Notably, CXCR4 transcript and protein was expressed by TC progenitors (TCPs), as well as TC II–IV (Figures 6D and S7E), while marginal zone FRCs were the dominant source of CXCL12, with higher expression at P14 relative to adults (Figures S2C, S7F and S7G), suggesting an early life CXCL12–CXCR4 axis may recruit TC progenitors to the subcapsular region, where local cues such as MRC-derived RANKL for TC I^25^, shape their differentiation.

To define the cellular environments of individual APC subsets, we performed neighborhood enrichment analyses separately for siLN and cLN at each age. CCR7⁺ cDC1 and cDC2 were enriched near multiple CD4 T cell populations and were prominent APC neighbors of naïve and proliferating CD4 T cells (Figures 6F–6I). However, within the T cell zone, these subsets occupied distinct cellular niches: CCR7⁺ cDC1 were preferentially associated with T_H_1 cells and proliferating or ISG^+^ T cell states and pDCs (Figures 6F and 6G), the latter a major source of type I IFN. In addition, CCR7⁺ cDC1 were enriched in proximity to CD8 T cells, consistent with their broader distribution extending into the CD8-rich deep T cell zone. Besides enrichment with naïve CD4 T cells, CCR7⁺ cDC2 exhibited a selective association with NKT cells (Figure 6H and 6I) which together with their preferential expression of CD1d (Figures S2D and S7H) suggests a role for CCR7^+^ cDC2 in presentation of microbiota-derived lipids and NKT cell activation.

By comparison, lymph-node resident cDC2A and cDC2B showed limited enrichment with conventional T cell states and instead occupied subcapsular/interfollicular niches with lymph-node specific patterns of interactions (Figure S7I). In peri-weaning siLN, cDC2A localized near MRCs, LTi cells, TC I and subcapsular sinus floor LECs, in line with previous reports of cDC2 uptake of lymph-borne antigens. In cLNs, both cDC2A and cDC2B subsets preferentially associated with HEVs. Resident cDC1 neighborhoods also changed with age: at P14, cDC1s were enriched near HEVs and BECs, whereas in adult lymph nodes their neighbors more frequently included CD4 and CD8 T cell populations (Figure S7J). Across ages, cDC1s retained proximity to NK cells and ILC1s, consistent with the cDC1–NK cell niche detected in our earlier Hotspot analysis (Figure 3A). The functional implications of these associations remain open, but the regional divergence illustrates how a common APC subset may encounter different cellular environments in anatomically and developmentally distinct nodes.

Given their marked enrichment during early life and essential role in pTreg priming, we next examined their cellular niche in peri-weaning nodes. Notably, naïve CD4 T cells and RORγt⁺ pTreg cells were among the most strongly enriched T cell neighbors of TC IV in peri-weaning cLNs (Figures 6J and K), revealing a specialized anatomical niche consistent with the established role of TC IV in directing pTreg cell differentiation.

Together, these findings reveal a striking spatial compartmentalization of specialized APC subsets within gut-draining lymph nodes. CCR7⁺ cDC1 occupied niches enriched for pDCs, T_H_1 cells and proliferating or ISG^hi^ T cell states, consistent with an interferon-responsive environment associated with effector T cell priming. By contrast, both CCR7^+^ cDC2 and tolerogenic TC IV localized alongside proliferating RORγt⁺ pTreg cells, while TC I occupied a distinct early-life subcapsular niche associated with MSM-like macrophages in contrast to the cDC2A/B enriched subcapsular niche in adults. This spatial organization provides a framework for understanding how the developing lymph node may coordinate antigen access and specialized APC–T cell interactions to shape the balance between intestinal tolerance and immunity.

### Intestinal TCs migrate to draining lymph nodes

The localization of TC IV within the T cell zone, alongside migratory CCR7⁺ cDC2s, raised the question of where these cells acquire intestinal antigens. Whereas lymph node-resident APCs can acquire soluble antigens delivered through afferent lymphatics and stromal conduits, migratory cDCs capture antigens within intestinal tissues before transporting them to draining lymph nodes. Given the established role of TC IV in priming pTreg cells specific for dietary and microbiota-derived antigens, we asked whether TCs similarly acquire antigens in the intestine before migrating to draining lymph nodes.

Studies of TCs have largely focused on gut lymph nodes, although our previous analysis of *Rorc^Venus^Prdm16^mBaojin^* reporter mice identified rare RORγt⁺CXCR6^−^PRDM16⁺MHCII^+^ cells within intestinal tissues^25^. To characterize these cells, we performed single-cell RNA sequencing (scRNA-seq) on CXCR6⁻RORγt(Venus)⁺MHCII⁺ cells isolated from the large intestine lamina propria (LI) and mLN of *Rorc^Venus^* P15–18 mice, using sample barcoding to retain their tissue identity. mLN-derived cells encompassed the four canonical TC subsets (TC I–IV), recently described TC progenitors (TCPs)^25^ , proliferating TCs and a contaminating population of CXCR6⁺ LTi cells (Figures 7A and S8A). While LI TCs were present within each of these clusters, the majority of LI TCs formed a cluster that was exclusively derived from intestinal tissue cells (Figures 7A and 7B). This cluster was heterogeneous, resolving into two distinct populations upon subclustering (Figure 7C). The first intestinal cluster contained cells expressing LTi- and TC progenitor-associated genes (*Cd34, Siglecf, Il1r2, Pdcd1, Zbtb16*) (Figures 7C and S8A), indicating that TCPs and LTiPs seed tissues as well as lymph nodes. The second cluster expressed genes associated with microbial sensing (*Clec10a, Mgl2, Ahr, Tlr2*), tissue residency (*Itgae, Hic1*) and retinoid uptake and signaling (*Stra6l, Rara*) (Figures 7C–E). Unexpectedly, these cells exhibited an epithelial-associated transcriptional program (*Epcam, Pigr*, *Krt8) including genes associated with epithelial cell adhesion (Jaml, Cdh1),* suggesting potential interactions with intestinal epithelial cells (IECs). Flow cytometric analysis confirmed the presence of PRDM16⁺ EpCAM⁺ TCs, co-expressing predicted markers including CLEC10A and CD103 (Figures 7F–H).

**Figure 7.**
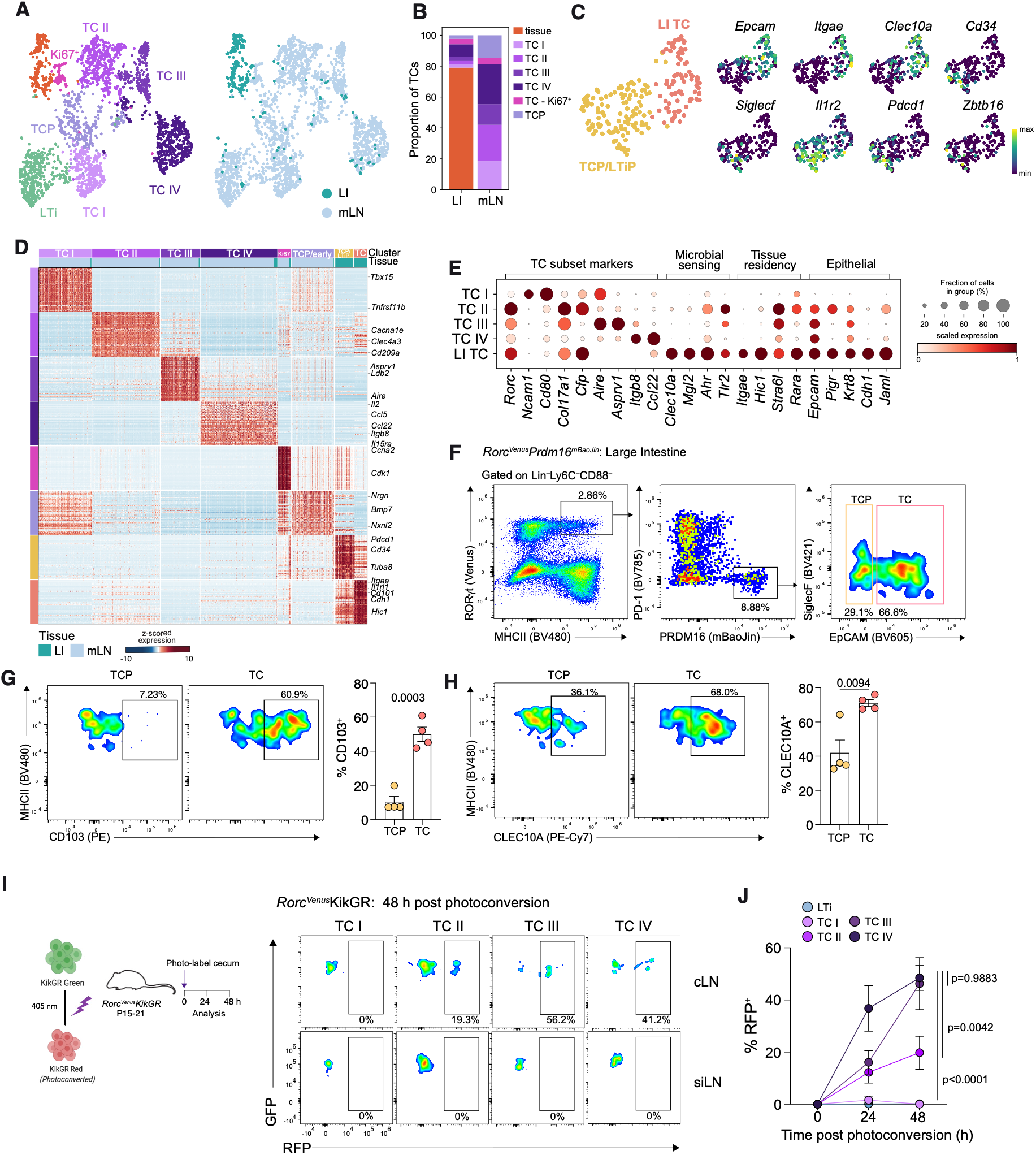
Intestinal Thetis cell populations exhibit distinct programs and migrate to draining lymph nodes. (A) UMAP of CXCR6^−^RORγt(Venus)^+^ TCs and LTi cells from large intestine and mLNs of P15-18 *Rorc^Venus^* mice profiled by scRNA-seq, colored by cluster annotation (left) and tissue of origin (right). (B) Proportion of each TC cluster among total cells from LI and mLN. (C) UMAP of subclustered LI cluster in (A), colored by subcluster (left) or expression of indicated genes (right). (D) Heat map showing scaled, imputed expression of top 50 DEGs (one versus the rest, fold change (FC) > log2(1.5), adjusted *P* < 0.01) for each mLN or LI Thetis cell cluster. (E) Dot plot with relative expression of selected DEGs in TC I-IV and LI TCs. (F) Representative flow cytometry of TCP and TCs in the large intestine lamina propria (LI) of P16 *Rorc^Venus^Prdm16^mBaoJin^* mice. (G) Representative flow cytometry of CD103 expression by TCP and TCs in the LI of P16 *Rorc^Venus^Prdm16^mBaoJin^*mice and summary graph (n = 4). (H) Representative flow cytometry of CLEC10A expression by TCP and TCs in the LI of P16 *Rorc^Venus^Prdm16^creERT^*^2^*^-mBaoJin^* mice and summary graph (n = 4). (I) Representative flow cytometry of RFP expression by TC subsets in cLNs and siLNs of P20 *Rorc^Venus^*KikGR mice 48 hr post-photoconversion. (J) Summary graph of RFP expression by TC subsets in cLNs of P15-21 *Rorc^Venus^*KikGR mice at indicated timepoints post-photoconversion (0 hr, n = 3; 24 hr, n = 8; 48 hr, n = 7). Data in F, G, H are representative of two independent experiments; data in I and J are pooled from five independent experiments. Error bars: mean ± s.e.m.; unpaired t-test (G and H) or two-way ANOVA (J). All P values are indicated on the corresponding graphs.

To further define their relationship to lymph node TCs, we applied single-cell hierarchical Poisson factorization (scHPF) to identify shared and subset-specific transcriptional programs. This identified distinct programs associated with TC I–IV, including a neural/adhesion program in TC I, a migratory program in TC IV comprising genes previously implicated in CCR7⁺ cDC migration, as well as a TC IV-specific Treg-inducing program (Figures S8B and S8C). Additional transcriptional modules were shared between TC progenitors and differentiated TC subsets. By contrast, intestinal TCs exhibited largely distinct transcriptional programs, suggesting adaptation to the intestinal tissue environment and potentially distinct functions within the tissue.

Given the localization of TC III and IV within the CCL19 rich T cell zone, their expression of CCR7, and expression of a migratory program by TC IV, we next investigated whether these subsets represented tissue TCs that migrate to the LN. To directly determine whether intestinal TCs migrate to draining lymph nodes, we used photoconvertible KikGR mice crossed to the *Rorc^Venus^* reporter and selectively photoconverted the cecum while shielding the cecal patch. Immediately following photoconversion, RFP^+^ cDCs and TCs were identified in the cecum but were undetectable in the cLNs, siLNs or cecal patch (Figure S9A). At 24 h, photoconverted cDCs were present among CCR7^+^ cDCs, selectively in cLNs, confirming regional lymphatic drainage and cDC migration (Figure S9B and S9C). Importantly, photoconverted TC III and IV were also detected in cLNs, increasing in frequency from 24 to 48 h (Figures 7I and 7J). A smaller fraction of TC II was labeled, whereas TC I remained unlabeled, consistent with their lack of CCR7 expression.

Together, these findings identify a transcriptionally distinct intestinal tissue TC population with features suggestive of microbial sensing and interactions with intestinal epithelial cells, alongside intestinal TCPs and rare TC I–IV cells. Photoconversion experiments establish tissue-to-node migration by TCs, with TC III and IV representing the predominant migratory subsets, whereas TC I and II appear largely lymph node resident. Whether migratory TC III and IV arise from intestinal TCPs, pre-existing differentiated TC III and IV, or the CD103⁺ intestinal TC population remains an important question for future investigation.

## DISCUSSION

Early-life encounters with dietary and microbiota-derived antigens preferentially induce pTreg cell responses, yet for many antigens, this capacity is restricted to a developmental window that closes with age. The decline in abundance of the pTreg-inducing TC IV population may contribute to this loss of tolerogenic capacity. However, an alternative, non-mutually exclusive possibility is that TC IV operates within a developmentally restricted cellular and anatomical environment that favors pTreg cell differentiation at the expense of alternative T cell fates. Our spatial atlas reveals profound differences in the tissue architecture, cellular composition, APC niches and cytokine environments in peri-weaning vs adult gut lymph nodes, with further specialization between small intestine- and colon-draining nodes. These findings suggest that the early-life window of intestinal tolerance may be determined not only by the availability of specialized tolerogenic TCs, but also by the spatial organization of the lymph node and the pathways through which intestinal antigens reach them.

One of the most striking features of peri-weaning lymph nodes was their distinct subcapsular compartment, enriched for developmentally restricted TC I, LTo-derived RANKL^+^ MRCs, and MSM-like sinus macrophages in place of classical SSMs. These MSM-like macrophages also occupied medullary sinuses but lacked CD209b, a key receptor for microbial polysaccharides, suggesting impaired complement dependent handling of microbial polysaccharides^55^. Regional differences were equally pronounced: cLNs exhibited a markedly reduced medullary sinus compartment compared with siLN, a distinction maintained into adulthood. Together, these findings suggest that developmental and regional specialization of the sinus network may influence the capture and distribution of lymph-borne antigens and their availability to distinct APC populations.

The replacement of classical antigen-relaying SSMs by an MSM-like population lacking expression of cell surface molecules that bind microbial antigens is particularly intriguing given the attenuated T_FH_ and germinal center responses observed in peri-weaning nodes. Reduced follicular immunity is a well-established feature of early life, attributed to delayed FDC maturation, incomplete T_FH_ differentiation and limited antigen availability, including suppression of microbiota-specific responses by maternally acquired antibodies^59,62–64^. Our findings suggest an additional anatomical checkpoint, whereby the early-life sinus macrophage network may restrict the delivery of intact antigens to follicular B cells and FDCs, consistent with the impaired antigen relay observed with SSM-to-MSM phenotypic diversion in response to altered RANK or LTβR signaling^57^. Intriguingly, TC I enrichment within the subcapsular niche, together with its expression of OPG and direct proximity to RANKL-expressing MRCs, raises the possibility that TC I contributes to this developmental organization. Whether TC I regulate macrophage differentiation, acquire lymph-borne antigens or perform other niche-organizing functions remains to be determined. The delayed emergence of follicular responses may reflect a developmental strategy that prioritizes tolerance induction while maternally acquired antibodies provide protection. Indeed, maternal antibodies actively suppress offspring T_FH_ and germinal center responses, with their absence accelerating endogenous intestinal IgA production^59,65^. Given previous evidence for reciprocal regulation between intestinal IgA responses and pTreg cells^8,66^, it is tempting to speculate that the developing subcapsular niche contributes to this temporal separation, favoring Treg cell priming while limiting competing follicular responses.

Indeed, in contrast to the limited T_FH_ cell response, peri-weaning lymph nodes were distinguished by widespread immune activation within the T cell zone, with particularly prominent CD4 T cell proliferation and pTreg differentiation in cLNs, contrasting with relatively quiescent T cell populations in adults. This activity occurred against a striking interferon-responsive transcriptional program spanning multiple immune and stromal lineages, preceding the previously described intestinal weaning reaction. Whether this program reflects type I or type II interferon signaling, the contribution of dietary or microbiota-derived stimuli, and how interferon signaling influences T cell priming remain important questions for future investigation. The reciprocal developmental regulation of IL-22 and its antagonist IL-22BP further highlights the distinct cytokine environments of young and adult nodes. Understanding how these developmental cytokine programs shape immune responses both locally and at distal intestinal sites will be important for defining how gut-draining lymph nodes coordinate intestinal tolerance and barrier immunity during early life.

Beyond these architectural and cytokine differences, our atlas reveals a spatial division of labor among specialized APC populations. CCR7⁺ cDC1 and cDC2 occupied distinct and overlapping territories within the T cell zone, with CCR7^+^ cDC1 preferentially associated with T_H_1 and CD8 T cell responses. In contrast, resident cDC1 occupied anatomically distinct niches that varied between siLNs and cLNs. TC subsets were similarly compartmentalized, with TC I and II concentrated in subcapsular and interfollicular regions, whereas TC III and IV localized to the T cell zone, where TC IV was engaged with naïve CD4 T cells and pTreg cells within the deep T cell zone. Notably, CCR7^+^ cDC2s shared the TC IV-pTreg niche despite previous genetic studies demonstrating that cDC antigen presentation is neither sufficient nor required for intestinal pTreg induction^9,15,16,18^. This raises the possibility that distinct molecular programs, rather than spatial positioning alone, enable TC IV to selectively direct pTreg differentiation. Defining these programs will be important for understanding the mechanisms underlying APC specialization and intestinal tolerance.

The timing of intestinal antigen exposure may provide an additional layer of regulation over the spatial organization of APC niches. At P14, we observed extensive CD4 T cell activation and pTreg differentiation in cLNs, whereas siLN exhibited comparatively limited responses. Intriguingly, this regional difference coincides with the developmentally regulated opening of goblet cell-associated antigen passages (GAPs), which emerge in the colon around P10 but remain largely absent from the small intestine until P18^67,68^. This temporal correspondence raises the possibility that epithelial antigen delivery is coordinated with the maturation of their lymph node niches, establishing distinct windows for tolerance to intestinal antigens. An important question is therefore how specialized APCs gain access to antigens within the intestinal tissue, and whether they exploit GAPs or alternative pathways of epithelial antigen transport.

Our identification of a transcriptionally distinct intestinal tissue TC population provides a starting point for addressing this question. Extending our previous characterization of the shared developmental pathway of LTi cells and TCs, we identified TC progenitors within the intestine, alongside PRDM16⁺EpCAM⁺ TCs expressing genes associated with microbial sensing, epithelial interactions and tissue adaptation. Although their precise anatomical localization remains unknown, their expression of epithelial-associated and adhesion molecules, including PIGR and CD103, suggests potential interactions with IECs that may facilitate antigen acquisition. Intriguingly, these transcriptional features are also present in human RORγt⁺ APCs (RORγt-DC-like cells), suggesting conservation of a tissue-associated TC program across species and potentially explaining their transcriptional divergence from the canonical TC subsets defined in mouse lymph nodes^26,69,70^.

We further demonstrate that intestinal TCs migrate to draining lymph nodes, identifying a potential route for delivering intestinal antigens to specialized APC niches within the T cell zone. Whether migratory TC III and IV differentiate from intestinal TC progenitors, intestinal TC III/IV counterparts, or the epithelial-associated TC population remains to be established. Equally important is understanding the signals that direct TC III versus TC IV differentiation, and whether these are acquired within intestinal tissues or following migration to the lymph node. Notably, although TC III also migrate from intestinal tissues, previous studies did not identify dietary antigen acquisition or pTreg-inducing activity by this subset^9^, raising the possibility that TC III and IV acquire distinct antigens and direct different T cell responses.

Together, these findings establish a spatial framework for understanding how the developing gut lymph node coordinates antigen access, APC specialization and T cell differentiation. By defining the anatomical organization of specialized APC niches, their developmental cytokine environments and the potential routes through which intestinal antigens reach them, our atlas provides a foundation for investigating how tolerogenic and immunogenic responses are partitioned during early life. Ultimately, defining how the timing and route of antigen exposure intersect with these developmentally regulated niches may inform recommendations for early-life food introduction to prevent allergy, as well as therapeutic strategies to restore tolerance in adulthood.

## METHODS

### Mice

*Rorc^Venus-creERT^*^2^ (also referred to as *Rorc^Venus^*), *Prdm16^creERT^*^2^*^-mBaojin^* (also referred to as *Prdm16^mBaojin^*) and KikGR mice have been previously described^15,25,71^. C57Bl/6 and *Ccr7^GFP^* mice were purchased from Jackson laboratories. Generation and treatments of mice were performed under protocol 21-05-007, approved by the Sloan Kettering Institute (SKI) Institutional Animal Care and Use Committee. All mouse strains were maintained in the SKI animal facility in specific pathogen free (SPF) conditions in accordance with institutional guidelines and ethical regulations. Both male and female mice were included in the study, and we did not observe sex-dependent effects. All mice analyzed were age and litter matched unless otherwise specified. The age of mice at time of analysis is included in the figures, legends and main text for individual experiments. No experimental interventions requiring randomization to control or experimental groups were performed in this study. No statistical calculations were performed to determine sample size.

### Xenium spatial transcriptomics

Mesenteric lymph nodes were collected from postnatal day 14 (P14) and adult (6 weeks old) C57Bl/6 mice (n = 3 for each age group). Samples were immersed in freshly prepared, ice-cold 10% formalin and fixed at 4°C overnight. After fixation, samples were washed twice in PBS for 30 min each at 4°C and dehydrated in 70% ethanol for 30 min at room temperature. Samples were stored in 70% ethanol at 4°C until paraffin processing. Samples were dehydrated through graded ethanol washes, cleared in Sub-X clearing agent, and infiltrated with paraffin. Samples were embedded using Leica HistoCore PEGASUS Parablocks and sectioned at 5 µm for downstream. Xenium in situ gene expression profiling was performed using the Xenium platform (10x Genomics) according to the manufacturer’s protocol. Sections were mounted on Xenium slides, deparaffinized, and subjected to target retrieval and protease treatment to enable probe accessibility. A customized 472-gene panel was used for probe hybridization, followed by ligation and rolling circle amplification to generate localized amplicons. Cyclic fluorescence imaging was performed on the Xenium Analyzer to detect RNA targets at subcellular resolution.

### Target panel design

A custom 472-gene Xenium panel was designed to resolve immune, stromal and APC subsets, cellular activation states and selected cytokine, chemokine and ligand–receptor pathways (**Table S1**). Panel design was performed using the 10x Genomics Xenium Designer in collaboration with the vendor. Reference single-cell datasets were used to assess anticipated gene expression and optical crowding across relevant cell populations. Highly abundant targets, including H2-Ab1, were excluded when predicted to impair accurate optical decoding.

### Spatial transcriptomic 10x Xenium data processing

Spatial transcriptomics was performed on tissue sections from postnatal day 14 (peri-weaning; n = 3 mice) and 6-week-old (adult; n = 3 mice) lymph node chains. Two adjacent sections were profiled using the Xenium platform (10x Genomics) with our custom panel. Each Xenium-defined image region comprised a tissue section from 3 peri-weaning or 3 adult mice, processed using Xenium 3-3.0.1.

### Transcript assignment and cell segmentation

Despite restricting analysis to transcripts overlapping clean DAPI-segmented nuclei, we observed significant contamination from transcript spillover in our segmented cells, identified by co-expression of two or more distinct lineage markers, likely due to transcript diffusion and z-plane contamination from overlapping cells. We therefore employed segger^40^, a graph-attention network method that leverages gene co-expression and spatial proximity to better assign transcripts to cells, and refine mapping of transcripts that overlap with neighboring cells due to diffusion or z-plane contamination. We used segger (version 0.2.0) with default parameters to assign transcripts to cells using DAPI staining from the Xenium Multi-Tissue Stain Mix, and ran the new transcript assignments through Xenium Ranger to create Xenium output bundles for each run. Segmentation quality was assessed by comparing annotations based on 10x nuclei assignments to segger based assigments, and the removal of contaminant genes.

### Pre-processing and quality control

Per-sample quality control metrics were computed and universal thresholds were set by inspection of metric distributions to filter low-quality cells for downstream analyses. Cells were required to have 20–300 counts and 10–100 genes to pass filtering. Further, cells with a segmentation area outside 5–200µm^2^ were excluded. Genes were required to be expressed in a minimum of 10 cells by raw counts to be included for further analyses. After ensuring that distributions of quality control metrics were consistent across all four Xenium samples, all cells retained after filtering were concatenated and analyzed together. This resulted in a final dataset of 1,267,407 cells and 470 genes. Cell counts were log1p transformed and normalized (‘log-normalized’), and then scaled without zero-centering. All data preprocessing was performed using SpatialData (v0.8.0) and scanpy (v1.12.1)^72,73^.

### Lineage assignment

Cells were assigned to broad lymphoid (T/NK/ILC), B/plasma cell, dendritic cell (pDC and cDC), Thetis cell, monocyte/macrophage, granulocyte or stromal lineage based on clustering using the a nearest-neighbor graph constructed with k=30 and cosine metric on log1p-normalized segger counts from all cells, and the top 50 principal components (PCs). Clustering was performed via Phenograph (resolution = 1.0)^74^, and clusters were manually assigned to broad cell type lineages using expression of marker genes for canonical immune or stromal cell subsets. Clusters that expressed marker genes from more than one lineage were labeled as a ‘mix’. Low-quality clusters with extremely low transcript counts were removed.

### Sublineage preprocessing pipeline

Within-lineage clusters were annotated based on differentially expressed genes, canonical lineage or subset-specific genes, reference transcriptomes and spatial localization within anatomically defined lymph node compartments. Reference annotations drew on consensus signatures from the Immunological Genome Project (ImmGen), and previously published single-cell datasets of Thetis cells (TCs), classical dendritic cells (DCs), lymph node stromal cells, LECs, and macrophages ^9,43,75–79^. Where analysis of clusters suggested further heterogeneity based on regional expression of lineage or subset-specific marker genes, selected clusters were further subclustered. Populations that co-expressed markers of two or more distinct cell lineages or cell types were annotated as ‘mixed’ cell types. These cells were excluded from subregion abundance analysis and were not included in data visualization plots, including spatial niche enrichment, lineage UMAPs and dot plots, and neighborhood enrichment plots. Low-quality cells, identified as subclusters with median counts per cell < 30, were removed from analysis.

### Cell type annotations

#### Dendritic cells

Dendritic cell annotations were informed by previously published dendritic cell single-cell atlases and subset-specific transcriptional signatures^75,76,80^. Among Flt3^+^ cDCs, cDC1s were distinguished by expression of *Xcr1*, *Tlr3* and *Naaa*, whereas cDC2s were annotated using *Sirpa*. Within the cDC2 lineage, subclustering resolved cDC2A and cDC2B, an interferon-stimulated gene (ISG)-high cDC2A population, based on established cDC2A- and cDC2B-specific transcriptional signatures. CCR7⁺ activated or migratory cDC1 and cDC2 populations were identified by expression of *Fscn1*, *Marcksl1* and *Apol7c*. CCR7^+^ cDC1 and CCR7^+^ cDC2 were distinguished by *Laptm4b*, *Gclc*, *Il12b* vs *Cd1d1*, *Il2ra* and *Fabp5* expression respectively.

#### Thetis cells

TC subsets were identified using previously defined TC I–IV signature genes ^9,15^ . A mixed cluster of *Apol7c^+^* TCs was subclustered to resolve TC III, TC IV and mixed TC/CCR7^+^ cDC populations.

#### T cells and innate lymphoid populations

αβ^+^ T cell populations (*Trbc1^+^Trbc2^+^*) were annotated using canonical marker genes for naïve, memory, activated, T helper and Treg cell subsets and cell states. Several transcriptionally heterogeneous clusters were further subclustered to resolve related populations. A cluster of Foxp3^+^ Treg cells was subclustered to distinguish RORγt⁺ peripheral regulatory T cell (pTreg)-associated and Helios⁺ thymic regulatory T cell (tTreg) populations distinguished by differential expression of *Rorc*, *Maf* and *Il10* (pTreg) vs *Ikzf2* expression (tTreg). Annotations were informed by published analyses of intestinal Treg cell heterogeneity^20,81–83^. A cluster of naïve T cells expressing *S1pr1*, was subclustered into CD4 and CD8 T cell populations. Naïve CD4 T cells were distinguished by *Cd4* and *Cd40lg* expression, whereas CD8 T cells were assigned based on absence of Cd4, along with expression of *Runx3*, *Ccr9* and *Itgae*^84^. Memory-like CD8 T cells were distinguished by *Eomes, Tbx21* and *Klra7* (Ly49G) expression^85,86^. A cluster of transcriptionally related innate-like lymphocyte populations was further subclustered to distinguish NKT cells (*Zbtb16*), γδ T cells (high *Trdc* and *Tcrg-C1*) distinguished by RORγt-and T-bet-associated transcriptional programs, and ILC2s (*Gata3*, *Il13* and *Klrg1*). A cluster of proliferating T cells (*Mki67^+^*) was further subclustered into proliferating CD4, CD8 and pTreg cell populations using their respective lineage signatures. A cluster of recently activated *Cd69*⁺*Nr4a1^+^* CD4 T cells was subclustered to resolve activated CD4, T_H_1 (*Tbx21*, *Ifng*), and Treg (*Foxp3^+^*) cells. In addition, we identified three minority clusters lacking known subset defining genes.

#### B cells and plasma cells

B cell annotations were informed by published transcriptional signatures of naïve, activated, germinal center and memory B cells. Plasma cells were identified using established plasma cell-associated transcriptional programs and further subclustered to distinguish IgA- and IgG-expressing populations based on *Igha* and *Ighg* constant-region transcripts, respectively.

#### Monocytes and macrophage

Monocyte/macrophage populations were annotated using canonical markers and published lymph node macrophage transcriptomic datasets, and their anatomical distribution within the subcapsular sinus, medullary sinuses, medullary cords and T cell zone^43,87^ .

#### Stromal and endothelial cells

Stromal annotations were informed by published lymph node stromal and lymphatic endothelial cell atlases^77–79^. Fibroblastic reticular cell (FRC) populations were distinguished using established transcriptional signatures of *Ccl19*^hi^ and *Ccl19*^lo^ T-zone reticular cells (TRCs), *Cxcl12*⁺ FRCs, marginal reticular cells (MRCs) and follicular dendritic cells (FDCs). MRCs were identified using *Tnfsf11* (RANKL) and other subset-associated markers, together with their localization beneath the subcapsular sinus. High endothelial venules (HEVs) were distinguished from other BECs by expression of *Ubd*. Lymphatic endothelial cells (LECs) were annotated using previously published single-cell atlases defining subcapsular sinus ceiling, floor and medullary sinus LEC populations. Ceiling LECs were identified using *Mmrn1* and *Cd36,* floor LECs by *Ccl20* and *Cd274*, and medullary sinus LECs by *Lyve1* and *Marco*, together with their anatomical localization. Two LEC clusters were further subclustered to segregate mixed cells from a population of medullary sinus LEC populations, and to further resolve a population of medullary sinus LECs, subcapsular sinus floor and ceiling LECs.

### Niche analysis

To define broad spatial niches, we computed the cell type composition of each cell’s neighborhood within a fixed radius of 40 µm. Neighbors were identified from spatial coordinates using a radius-based nearest-neighbor search in scikit-learn (v1.7.2), restricted to cells from the same tissue section and to cells with “clean” cell type annotations; the cell itself was excluded. Neighborhood composition was calculated as the fraction of neighbors belonging to each fine cell type annotation, and the number of valid neighbors was recorded per cell. Cells with fewer than 6 valid neighbors were excluded from niche analysis leaving 1,192,240 of 1,200,309 cells (99.3%). Neighborhood fractions were standardized (z-scored per cell type) and clustered using k-means on scikit-learn (v1.7.2) with random_state=0, n_init=’auto’. We evaluated k = 2-9 using within-cluster inertia and mean silhouette score, and selected k = 6 for cluster stability and biological interpretability. Differences in niche proportions between age groups were tested using each animal as a biological replicate with propeller (speckle, v1.12.0)^88^, which applies a moderated t-test (limma) to logit-transformed sample-level proportions, with Benjamini-Hochberg correction. Excluded cells were left unassigned.

### Anatomical region assignment

For each Xenium sample, cLN and siLN subregions were assigned using the orientation and drainage anatomy of the mesenteric lymph node chain at the time of embedding, and subsequently visualized on Xenium Explorer (v4.1.1). Each siLN chain was further segmented into four regions draining proximal to distal small intestine (S1 (proximal) to S4 (distal)), using MRCs and SCS LECs to guide boundaries between individual nodes. Concordance between biological and adjacent section replicates for each subregion were determined by comparing broad cell type composition and QC metrics including counts per cell, genes per cell, and number of neighbors in a fixed radius. One adult siLN biological replicate was excluded from downstream regional and neighborhood analyses due to low cell density and low number of genes per cell. For one peri-weaning siLN chain and one adult cLN, the tissue section resulted in a partial capture of the lymph node or chain that was not representative of a longitudinal section through the meridian. Since cell type composition could not be directly compared between these tissue regions, these samples were excluded from differential gene analysis and analyses of subregion cell type abundance.

For all remaining replicates and subregions, we assessed global transcriptional similarity using scikit-learn (v1.7.2). Raw counts were summed across all cells within each sample-chain unit and size factors were estimated PyDESeq2 (v0.5.4) using the default median-of-ratios method^89^. Counts were variance-stabilized with PyDESeq2 agnostic of sample labels, VST values were mean-centered per gene (without scaling), and principal component analysis was performed.

### Differential gene expression analysis

To identify age-associated transcriptional differences, we performed pseudobulk differential expression analysis, both globally and per-cell type. Raw UMI counts were summed across all cells from each animal with decoupleR (v2.2.0)^90^, producing pseudobulk expression profiles per animal and thereby merging adjacent sections and nodes from the same animal. Only pseudobulk profiles comprising at least 10 cells and at least 100 total counts were retained. Genes were tested only if they had at least 10 counts in at least 2 profiles. Differential expression between Young and Adult animals was tested with PyDESeq2 (v0.5.4) using default median-of-ratios size-factor normalization and dispersion estimation, with a negative binomial model and the design ∼ frac_siLN + Age. Because the proportion of siLN versus cLN cells differed between animals and age groups, the fraction of each animal’s cells originating from siLN (frac_siLN) was included as a covariate to account for regional composition. Wald test *P*-values were adjusted for multiple testing using the Benjamini-Hochberg procedure, and genes with an adjusted *P* -value < 0.05 were considered differentially expressed.

To compare gene expression across ages and gut-draining LN regions, pseudobulk profiles were generated for each animal and region (siLN or cLN), retaining profiles with at least 10 cells, 100 total counts per profile and genes with at least 10 counts in two or more profiles. Differential expression was assessed using PyDESeq2 with the design ∼ group. Four pairwise comparisons were evaluated using Wald tests: peri-weaning vs adult within siLN, peri-weaning vs adult within cLN, and cLN vs siLN within each age group. *P*-values were adjusted for multiple-testing using the Benjamini-Hochberg method and genes with *P* < 0.05 were considered significant.

### Gene signature scores

To compare interferon responses across cell types and ages, we defined an interferon-stimulated gene (ISG) signature by intersecting an ISG geneset, filtered on genes within our Xenium dataset, with genes differentially expressed across ages or lymph node regions. Per-cell ISG scores were calculated from log-normalized expression data using scanpy.tl.score_genes and scaled across all cells.

### Receptor Ligand co-localization analysis

To compare ligand–receptor (LR) activity across spatial niches, we used the bivariate spatial metrics implemented in LIANA+ (v1.8.1)^91^. The low-density region in A3 siLN and cells left unassigned to a niche were excluded. A spatial connectivity graph was built from cell coordinates using a Gaussian kernel (40 µm bandwidth), consistent with the spatial scale used for niche definition. Edge weights below 0.1 were set to zero, and self-connections were excluded. Bivariate scores were calculated from log-normalized expression data for LR pairs in the CellPhoneDB database implemented in LIANA+, using cosine similarity as the local metric and Moran’s R as the global metric (100 permutations, retaining negative values)^92^ . Only LR pairs expressed in at least 0.1% of cells were evaluated. Local scores were averaged across cells within each niche and z-scored for each LR pair across niches to identify relative spatial enrichment independent of absolute score magnitude.

To compare cell–cell communication networks across ages, we performed separate analyses of peri-weaning and adult siLNs using LIANA+. For each age group, interactions between fine cell types were inferred using rank_aggregate with the mouse CellPhoneDB database. Interactions were scored using log-normalized expression data, requiring expression in at least 10% of cells within a cell type (default threshold), and weighted by spatial proximity using a Gaussian connectivity graph (40 µm bandwidth).

For visualization, interactions were restricted to cell types retained within each age group and LR pairs with a CellPhoneDB permutation *P* < 0.05, for which both ligand and receptor genes were included in the gene set of interest. Interaction strength was represented by CellPhoneDB lr_means, and self-interactions were excluded. Age-specific communication networks were visualized as directed multigraphs using NetworkX (v3.5), with edge widths proportional to the square root of lr_means and node sizes proportional to the square root of the number of cells belonging to each cell type.

### Neighborhood enrichment analysis

Spatial co-occurrence of cell types was assessed using neighborhood enrichment analysis in Squidpy (v1.8.1)^93^. Spatial neighbor graphs were constructed from cell coordinates using a fixed radius of 20 µm (non-Delaunay, generic coordinates), separately for each tissue section (Animal_ID_sub) to prevent connections between sections. Low-density regions and cells lacking a subregion (siLN/cLN) assignment were excluded from analysis. Neighborhood density was assessed by examining the mean number of neighbors per cell in each sample.

Neighborhood enrichment was calculated separately for each age–region combination, excluding cell types represented by fewer than 10 cells. Groups containing fewer than two cell types or no spatial connections were excluded. For each cell type pair, the observed number of neighboring connections was compared with a null distribution generated by 1,000 permutations of cell type labels (random seed = 0), and enrichment was expressed as a z-score. Infinite z-scores arising from zero variance in the null distribution were treated as missing values. Neighborhood enrichment z-scores were visualized across ages or regions and the top enriched cell types were plotted. For spatial overlays of cell type interaction neighborhoods, we overlaid fine cell type annotations on the default cell segmentation masks from Xenium 3-3.0.1 and Xenium Ranger using the Multi-Tissue Stain Mix.

### Macrophage subset gene signatures

Macrophage subset signatures were derived from published bulk RNA-seq data on FACS-isolated intestinal lymph node macrophage populations (GSE282129)^43^. Using the provided counts per million (CPM) values and sample metadata, samples from small intestinal lymph nodes were included, and genes with CPM ≥ 1 in at least three samples were retained (18,818 of 21,750 genes). Expression values were log2(CPM + 1) transformed, and differential expression between macrophage subsets (SSM, MSM, and MCM) was assessed using ordinary least squares linear regression, with macrophage subset as the variable of interest and tissue site as a covariate (statsmodels v0.14.6). Pairwise differences between subsets were evaluated using t-tests. Subset-specific signatures were defined as genes significantly upregulated relative to both other subsets (log2 fold change ≥ 0.5, adjusted *P* < 0.05 in both comparisons), with mean CPM ≥ 10 in the corresponding subset. Genes were filtered for genes represented in our Xenium gene panel, yielding signatures comprising 48 SSM, 21 MSM, and 63 MCM genes. Per-cell signature scores were calculated from log-normalized expression data using scanpy.tl.score_genes.

### Cell fate analysis

Macrophage differentiation trajectories were inferred using Palantir (v1.4.5)^94^. A diffusion map was constructed from the top 20 principal components using 20 nearest neighbors (k = 20). The multiscale diffusion space was then computed using Palantir’s default eigengap-based selection, retaining two diffusion components (DC1 and DC2). The Mac-1 cell with the lowest DC2 value, located at one extreme of the diffusion space, was selected as the starting cell. Palantir was run using 1,200 waypoints and 30 nearest neighbors (k = 30) to infer pseudotime, differentiation potential (entropy), and terminal-state fate probabilities.

### Spatial proximity analysis

To compare the spatial proximity of macrophage populations to the subcapsular sinus floor across ages, we calculated the Euclidean distance (µm) from each source cell to its nearest SCS fLEC within the same tissue section. Nearest-neighbor distances were computed from two-dimensional spatial coordinates using k-d tree queries (scipy.spatial.cKDTree, SciPy v1.18.0)^9^. Analysis was restricted to peri-weaning and adult siLN, given paucity of medullary sinus macrophages in cLNs, with SCS cLECs, Mac - 1, and Mac - 2 as source populations in peri-weaning mice, and SCS cLECs, Mac - SCS, and Mac - 3 in adults.

To account for the greater abundance of SCS fLECs in peri-weaning lymph nodes, which could bias nearest-neighbor distances toward shorter values, we repeated the analysis after downsampling SCS fLECs to match adult abundance. Within each tissue section, a fixed number of SCS fLECs was randomly sampled without replacement, and distances from each source cell to its nearest sampled SCS fLEC were calculated. This procedure was repeated 100 times (random seed = 0), and distances were averaged across iterations. Sections containing fewer SCS fLECs than the required sampling number or fewer than five source cells were excluded. Median distances were then calculated for each source population within each tissue section for comparison across ages.

### Spatial transcriptomic 10x Visium HD analysis

Lymph node sections adjacent to those used for Xenium analysis were placed on the capture area of Visium Spatial Gene Expression for FFPE slides (10x Genomics). Slides were placed in a drying rack, incubated at 42°C for 3 h and then transferred to a desiccator overnight at room temperature. Slides were subsequently processed according to the 10x Genomics Visium FFPE workflow. Tissue sections were deparaffinized and stained with hematoxylin and eosin (H&E) to visualize tissue morphology. Library quality was assessed using the Agilent Bioanalyzer High Sensitivity DNA Kit (Agilent Technologies, 5067-4626). Libraries were sequenced on an Illumina NovaSeq X Plus platform using paired-end sequencing. Sequencing was performed with the following configuration: 28 bp for Read 1, containing the spatial barcode and UMI; 10 bp each for the i7 and i5 indexes; and 50 bp for Read 2, containing transcript sequences.

### Visium HD Data preprocessing and Quality Control

Samples were processed using Space Ranger (v4.0.1), and expression data binned at 8 µm resolution were used for downstream analyses. Quality control metrics, including total counts, number of detected genes, and proportions of mitochondrial and hemoglobin transcripts, were calculated for each bin. Bins were retained if they contained 60–800 counts, 50–700 detected genes, <4% mitochondrial and <1% hemoglobin counts. An additional 463 bins corresponding to artifacts or outlier tissue regions identified by manual inspection in Loupe Browser were excluded. Genes not detected in any remaining bins were removed, yielding a final dataset of 407,338 bins and 19,045 genes. Following quality control, raw counts were library-size normalized, log-transformed, and scaled. Data preprocessing was performed using SpatialData (v0.8.0) and scanpy (v1.12.1).

### Spatial Hotspot analysis

Spatially co-varying gene expression programs were identified in 8 µm-binned Visium data using Hotspot (v1.1.3)^46^ . Analysis was performed on raw counts from quality-controlled bins using a Bernoulli (presence/absence) model, with total UMI counts per bin as a covariate. A spatial k-nearest-neighbor graph was constructed from bin coordinates (k = 60, unweighted), and spatial autocorrelation was calculated for each gene. Genes exhibiting significant spatial autocorrelation (FDR < 0.05) were retained for pairwise local correlation analysis and grouped into modules using Hotspot’s agglomerative clustering algorithm (FDR threshold = 0.05; minimum module size = 30 genes). Genes not assigned to a module were excluded from downstream module analyses. Spatial autocorrelation was computed on CPU, while pairwise local correlations were calculated using GPU acceleration with CuPy (v14.1.1). Per-bin module scores were calculated using calculate_module_scores and visualized spatially to compare expression programs across ages.

### Tissue processing for flow cytometry and scRNA-seq

Mice were euthanized by CO2 inhalation. Organs were collected and processed as follows. Lymphoid organs were digested in collagenase in RPMI1640 supplemented with 5% fetal calf serum, 1% L-glutamine, 1% penicillin–streptomycin, 10 mM HEPES, 1 mg ml^−1^ collagenase A (Sigma,11088793001) and 1 U ml^−1^ DNase I (Sigma, 10104159001) for 45 min at 37 °C, 250 rpm. The small and large intestines were harvested, and Peyer’s patches were removed from the small intestines while cecal patches were removed from the large intestines. The intestines were flushed with PBS and incubated in PBS supplemented with 5% fetal calf serum, 1% L-glutamine, 1% penicillin–streptomycin, 10 mM HEPES, 1 mM dithiothreitol, and 1 mM EDTA for 15 min at 37 °C, 250 rpm, to remove the epithelial layer. Samples were washed and incubated in digest solution with ceramic beads (0.25 inch) (MP Biomedicals, 116540034) (3 per sample) for 30 min at 37 °C, 250 rpm for tissue dissociation. Digested samples were filtered through 100-µm strainers and centrifuged to remove collagenase solution.

### Flow cytometry

For flow-cytometric analysis, dead cells were excluded by staining with LIVE/DEAD Fixable Zombie NIR in PBS for 10 min at 4 °C, with anti-CD16/32 to block binding to Fc receptors before cell-surface staining.

Extracellular antigens were stained for 30 min at room temperature in staining buffer (2% FBS, 0.1% Na azide, in PBS), diluted 1:1 with Brilliant Violet (BD Biosciences) staining buffer. For intracellular protein analysis, cells were fixed and permeabilized with Cytofix (BD Biosciences) and/or an eBioscience Foxp3 kit, as per the manufacturer’s instructions. Intracellular antigens were stained for 30 min or overnight at 4 °C in the respective 1× Perm/Wash buffer. The antibodies used for flow cytometry and FACS are listed in **Table S2**. Unless otherwise stated, we used the following gatings: TCs, Lin (Siglec-F^hi^, TCRβ, TCRγδ, CD19, B220, NK1.1, CD88, Ly6C/6G)^−^RORγt^+^CXCR6^−^MHCII^+^; LTis, Lin^−^RORγt^+^CXCR6^+^CCR6^+^MHCII^+^; cDC1s, Lin^−^ RORγt^−^CD11c^+^MHCII^+^CD11b^−^XCR1^+^; cDC2s, Lin^−^RORγt^−^CD11c^+^MHCII^+^CD11b^+^XCR1^−^. Samples were acquired on a Cytek Aurora. FACS isolation was done using a BD Aria.

### Isolation of TCs for 10x Genomics Flex gene-expression profiling

The mLN and large intestine from P15–18 *Rorc^Venus^* mice (n = 67 for large intestine and n = 22 for mLN), were processed as outlined above. Single cell suspensions were depleted of lineage (TCRβ, TCRγδ, CD19, B220, NK1.1, Ly-6G)^+^ cells by staining with biotinylated antibodies followed by magnetic bead (Dynabeads M-280 streptavidin; Invitrogen) negative selection. Cells were incubated with anti-CD16/32, and extracellular antigens were stained for 30 min at room temperature in sorting buffer (2% FBS, 2 mM EDTA, in PBS). Cells were washed and resuspended in cRPMI with SYTOX blue (Invitrogen) to exclude dead cells.

### Single-cell RNA sequencing using the 10x Genomics Flex workflow

Lin^−^RORγt(Venus)^+^CXCR6^−^MHCII^+^ cells were sorted into 500 µl of fixation buffer (4% formaldehyde, 1× Conc. Fix and Perm Buffer B; 10x Genomics, PN-2001301) and processed according to the 10x Genomics protocol for fixation of cells and nuclei. Briefly, cells were fixed for 16–24 h at 4 °C. To terminate fixation, cells were centrifuged at 850 × g for 5 min at room temperature and resuspended in 1 mL of Quenching Buffer (1× Quench Buffer; 10x Genomics, PN-2000516). Preparations were subsequently supplemented with 0.1 volumes of Enhancer (10x Genomics, PN-2000482) and 10% glycerol for long-term storage. Samples were thawed at room temperature, centrifuged at 850 × g for 5 min, and resuspended in 1 mL of 0.5× PBS containing 0.02% BSA. Hybridization reactions were prepared in 80 µL containing 20 µL of Mouse Whole Transcriptome Analysis (WTA) probes (10x Genomics, PN-2000718) and incubated at 42 °C for 16–24 h according to the manufacturer’s recommendations. Following hybridization, samples were diluted in Post-Hybridization Wash Buffer. Cells from individual hybridization reactions were pooled and washed three times in Post-Hybridization Wash Buffer for 10 min at 42 °C, resuspended in Post-Hybridization Resuspension Buffer, filtered through a 30-µm Miltenyi Biotec filter. GEM generation was performed according to the 10x Genomics protocol, where samples were loaded onto a Chromium Chip Q and processed on a Chromium X instrument (10x Genomics). Following GEM generation, GEMs were recovered and processed according to the manufacturer’s instructions, and the resulting products were pre-amplified and indexed to generate sequencing libraries. Final libraries were sequenced on an Illumina NovaSeq X using standard dual indexing and demultiplexing. Raw BCL files were processed using Cell Ranger (10x Genomics).

### Single-cell RNA-seq processing and annotation

Samples were profiled with the 10x Genomics GEM-X Flex v2 Gene Expression assay (Chromium Mouse Transcriptome Probe Set v2.0.0, 19,070 targeted genes) and pooled across six probe barcodes (one mLN and five large intestine samples). FASTQ files were processed with Cell Ranger v10.1.0 multi against the GRCm39-2024-A mouse reference to generate per-sample filtered count matrices. Per-sample matrices were loaded with scanpy (v1.12.1) and per-cell quality control metrics were computed. Cells were filtered based on the number of RNA-seq transcripts (<1,808 or >35,000 counts), the number of detected genes (>1,339 and <8,103 detected genes, mitochondrial counts (<4%) and hemoglobin counts (<1%), leaving 3,091 of 3,660 cells. Finally, mitochondrial, hemoglobin, Malat1 and genes detected in <2 cells were discarded, leaving 15,440 genes. Cluster labels

### Dimensionality reduction, cell clustering, and visualization

The filtered count matrix was library-size normalized, log-transformed (‘log-normalized’ expression values) and then centered and scaled (‘scaled’ expression values) using scanpy (v1.12.1). Cells were clustered using PhenoGraph (v1.5.7) using a nearest-neighbor graph constructed with k=20, resolution=0.3 on the top 30 PCs, yielding 14 clusters. Clusters were manually annotated based on expression of canonical TC I-IV, TCP, LTiP, LTi genes^9,25^. Minority contaminating clusters of stromal cells (*Ptprc*^−^), monocyte/macrophages and a cluster of unknown identity (*Rorc*^−^) were excluded from downstream analysis. To resolve heterogeneity within a cluster of intestinal TCs, principal component analysis (PCA) was repeated on this subset, retaining the top 20 principal components to construct a k-nearest-neighbor graph (k = 15, Euclidean distance). Cells were then clustered using the Leiden algorithm (resolution = 0.3, run to convergence) and the resulting two subclusters were visualized on the original UMAP embedding and annotated based on TC and LTi subset signature scores.

### Differential gene expression tests

Differentially expressed genes (DEGs) between groups of cells were identified in Seurat (v6.0.0) using MAST (MAST v1.28.0)^95^ via FindMarkers. Raw counts, log-normalized expression, UMAP coordinates, and metadata were transferred from the scanpy object into a Seurat object using anndata (v0.8.0). DEGs were identified in one-vs-rest tests comparing each cluster against all other retained cells. To account for differences in detection rate, the scaled number of genes detected per cell (cellular detection rate) was included as a latent variable. DEGs were reported according to their log-fold change (>1.5) and adjusted *P*-value (<0.01).

### Data imputation for scRNA-seq data

MAGIC (v3.0.0) imputation was applied to the log-normalized expression values to further de-noise and recover missing values^96^ . Imputed gene expression values were used for data visualization only on heat maps, where stated.

### Gene signature scores

TC signature scores were computed using Smart-seq3 (SS3) reference dataset of TC and progenitor populations generated in Paucar Iza et al. (GSE316677)^25^. Marker genes for each TC and LTi cell population were identified by one-versus-rest differential expression with MAST as described above and filtered to genes upregulated in the population of interest (adjusted *P* < 0.01, fold change > 1.5). Signature scores for each population were calculated from log-normalized expression data using the corresponding marker gene sets and scanpy.tl.score_genes with default parameters.

### scHPF factorization

Gene expression programs were identified using single-cell hierarchical Poisson factorization (scHPF, v0.5.0) to raw UMI counts from all TC subset and progenitor clusters^97^. Genes detected in fewer than 10 cells were removed, yielding a matrix of 2,117 cells and 12,288 genes. Five independently initialized models were trained for each K (8 to 13), and the trial with the lowest final negative log-likelihood was retained. K=13 was selected based on seed reproducibility and biological interpretability. Per-cell factor scores were normalized to sum to one across factors.

### Photoconversion surgery

P15-21 KikGR mice were anesthetized with isoflurane [1–3% by continuous inhalation], and given Buprenorphine (Ethiqa XR, 3.25 mg/kg) and Meloxicam (2 mg/kg) by subcutaneous injection for pain relief during surgery. Eye lubricant was applied, and the fur around the abdomen was shaved. Mice were placed on a heating pad in a supine position. The abdominal skin was disinfected with 70% ethanol and povidone-iodine before surgery. A ∼1 cm vertical incision was made in the skin on the lower abdomen, and Bupivacaine was administered to the exposed peritoneal tissue for local anesthesia. A ∼0.5-cm incision was made in the abdominal wall, through which the cecum was externalized gently. A piece of sterile aluminum foil was arranged on either side of the cecum and covered the tip of the cecum, shielding the skin, abdominal cavity, and cecal patch. Photoconversion was achieved by illuminating the cecum with a 405-nm handheld light source (Prizmatix) positioned approximately 5 cm above the tissue. The light source was applied to the cecum for 2.5 min per side on both sides while phosphate-buffered saline was applied to rinse and maintain hydration. After photoconversion, the cecum was gently replaced, and the abdominal cavity and skin were closed separately with sutures (6–0 Vicryl). Mice were allowed to recover under a heating lamp. Mice were analyzed at the indicated timepoints post surgery or immediately after photoconversion to assess photoconversion spillover to lymph nodes.

### Quantification and statistical analysis

Analysis of all data was done using unpaired two-tailed t-tests, one- or two-way ANOVA with a 95% confidence interval, or model-based analysis of single-cell transcriptomics, as specified in the text or figure legends. Statistical tests for flow cytometry data were done with GraphPad Prism v.11. *P* < 0.05 was considered significant: * *P* < 0.05; ** *P* < 0.01; *** *P* < 0.001; **** *P* < 0.0001. Details of the number of replicates, sample size, significance tests, the value and meaning of n for each experiment and exact *P* values are included in the Methods or figure legends. The Flex scRNA-seq experiment was carried out once. No measures were taken to estimate sample size or to determine whether the data met the assumptions of the statistical approaches used.

## ACKNOWLEDGMENTS

We thank Yuri Pritykin, Amanda Sun and Tamjeed Azad for helpful discussions; Theresa Lu for KikGR mice; Ronan Chaligne, Catherine Snopkowski and Karolis Kumpaitis (Single-cell Analytics and Innovation Lab (SAIL)); Ning Fan (Molecular Cytology); and the Integrated Genomics Operation core facilities at Memorial Sloan Kettering Cancer Center for technical assistance.

## AUTHOR CONTRIBUTIONS

Conceptualization, Y.L. and C.C.B.; investigation, Y.L., G.F., A.B., Y.P., Y.A.P.I., G.S., T.H. and V.C.; formal analysis, Y.L. and Y.A.P.I.; bioinformatics, P.L., A.Y. and T.P.; writing – original draft, Y.L., C.C.B. and P.L.; writing – review and editing, Y.L., C.C.B. and P.L.; supervision and funding acquisition, C.C.B..

## FUNDING

This work was supported by a NIH NIAID DP2 award (DP2AI171116) (C.C.B.), Pew Biomedical Scholar Award (C.C.B), the Mathers Foundation, and a NCI Cancer Center Support Grant P30 CA08748. C.C.B. is a Freeman Hrabowski Scholar, Howard Hughes Medical Institute (HHMI). Y.A.P.I. is supported by an HHMI Gilliam Fellowship. G.S. is supported by a Kravis-WiSE fellowship.

## DECLARATION OF INTERESTS

The authors declare no competing interests.

## DATA AND CODE AVAILABILITY

All data needed to evaluate the conclusions in the paper are present in the paper or the Supplementary Materials. No new software or code was generated in this study. The scripts used to analyze spatial transcriptomics data can be accessed at Github (https://github.com/The-Brown-Lab-MSKCC/Spatial_mLN).The sequencing data are available through the Gene Expression Omnibus under accession GSE (pending). Further information and requests for materials may be directed to, and will be fulfilled by, corresponding author C.C.B..

## SUPPLEMENTARY FIGURES

**Supplementary Figure 1.**
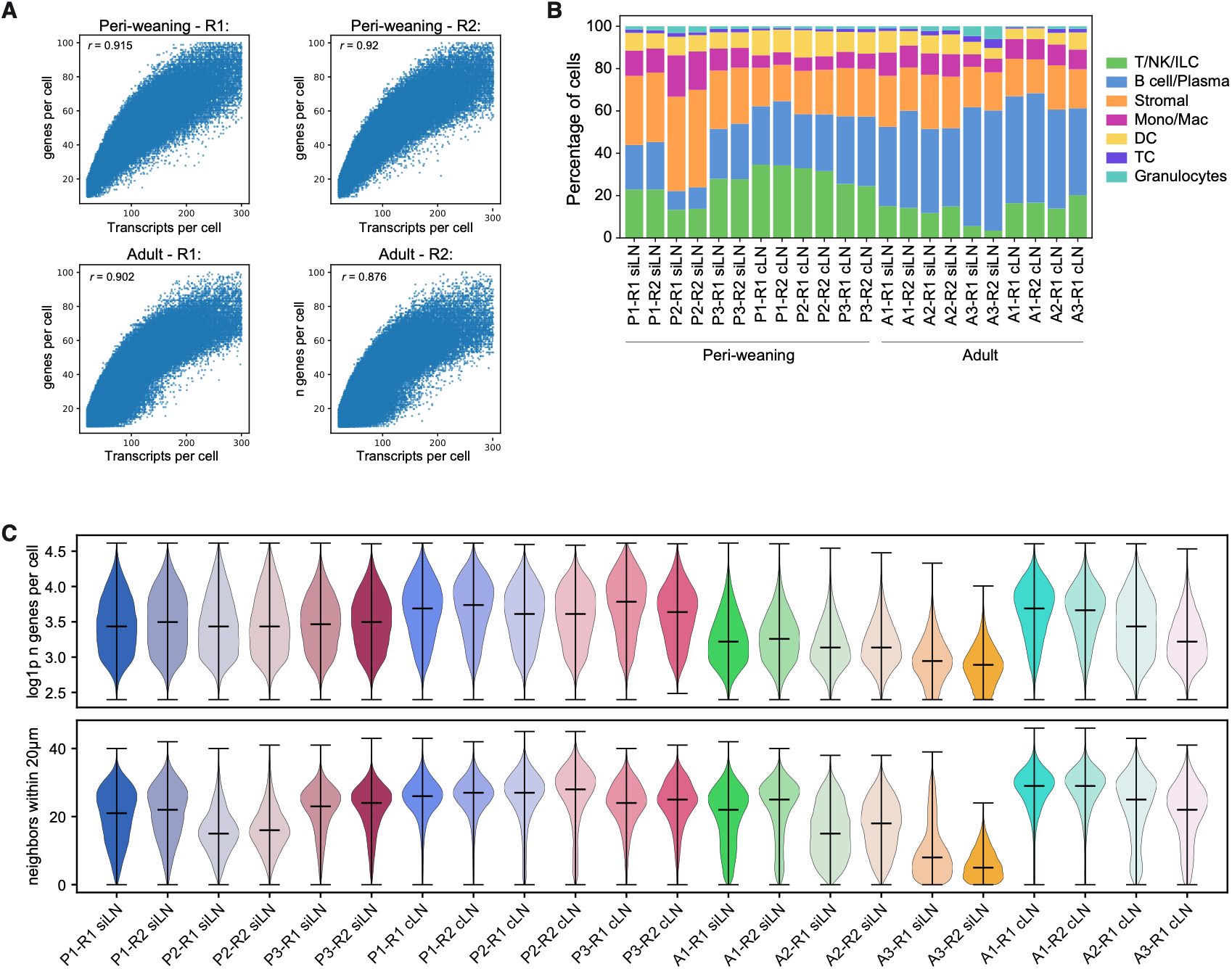
Xenium profiling quality control. (A) Number of transcripts (x) and genes (y) detected per cell within each Xenium-defined image region (R). Each dot represents one cell. *r* = Pearson correlation coefficients. (B) Cell type composition of each sample. Samples labeled by animal (P1-P3, peri-weaning; A1-A3, adult), Xenium-defined region (R1, R2), and anatomical region (siLN, cLN). (C) Number of detected genes per cell (top) and number of neighboring cells within 20 µm (bottom) for each sample. Horizontal lines indicate the median. A3-siLNs were excluded from downstream analysis due to low QC.

**Supplementary Figure 2.**
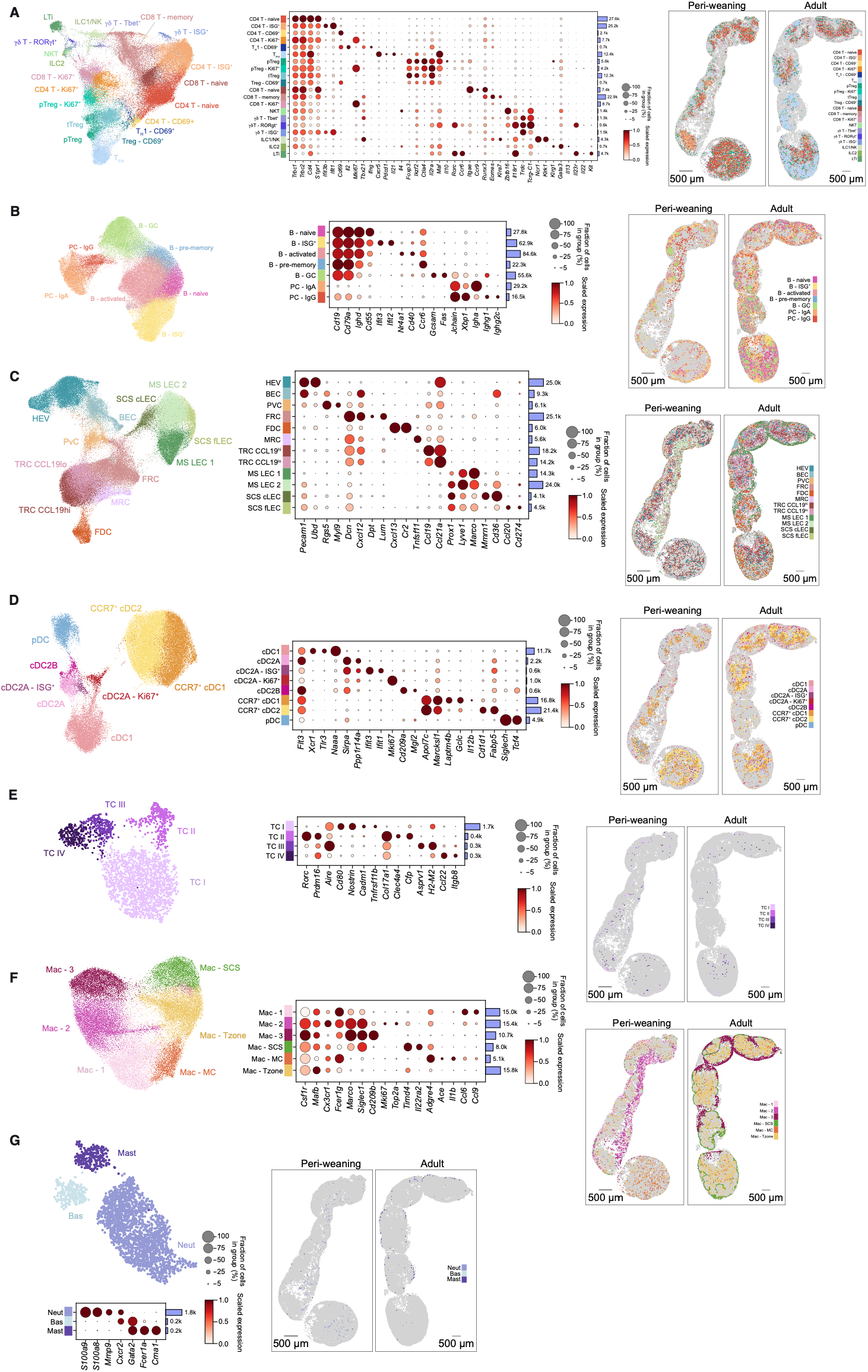
Spatially resolved single-cell annotation resolves diverse immune and stromal populations. (A) – (G) Subclustering of cells within each broad lineage: (A) T/NK/ILC, (B) B cell/plasma, (C) stromal, (D) dendritic cell (DC), (E) Thetis cell (TC), (F) monocyte/macrophage, and (G) granulocyte. For each lineage, UMAP embedding is colored by fine cell type. Dot plot with relative expression of selected marker genes for each fine cell type. Bar plots to the right of each dot plot indicate the number of cells in each fine cell type cluster across peri-weaning and adult lymph nodes. Spatial overlays show the location of fine cell types in representative peri-weaning and adult lymph node chains. Scale bars, 500 µm.

**Supplementary Figure 3.**
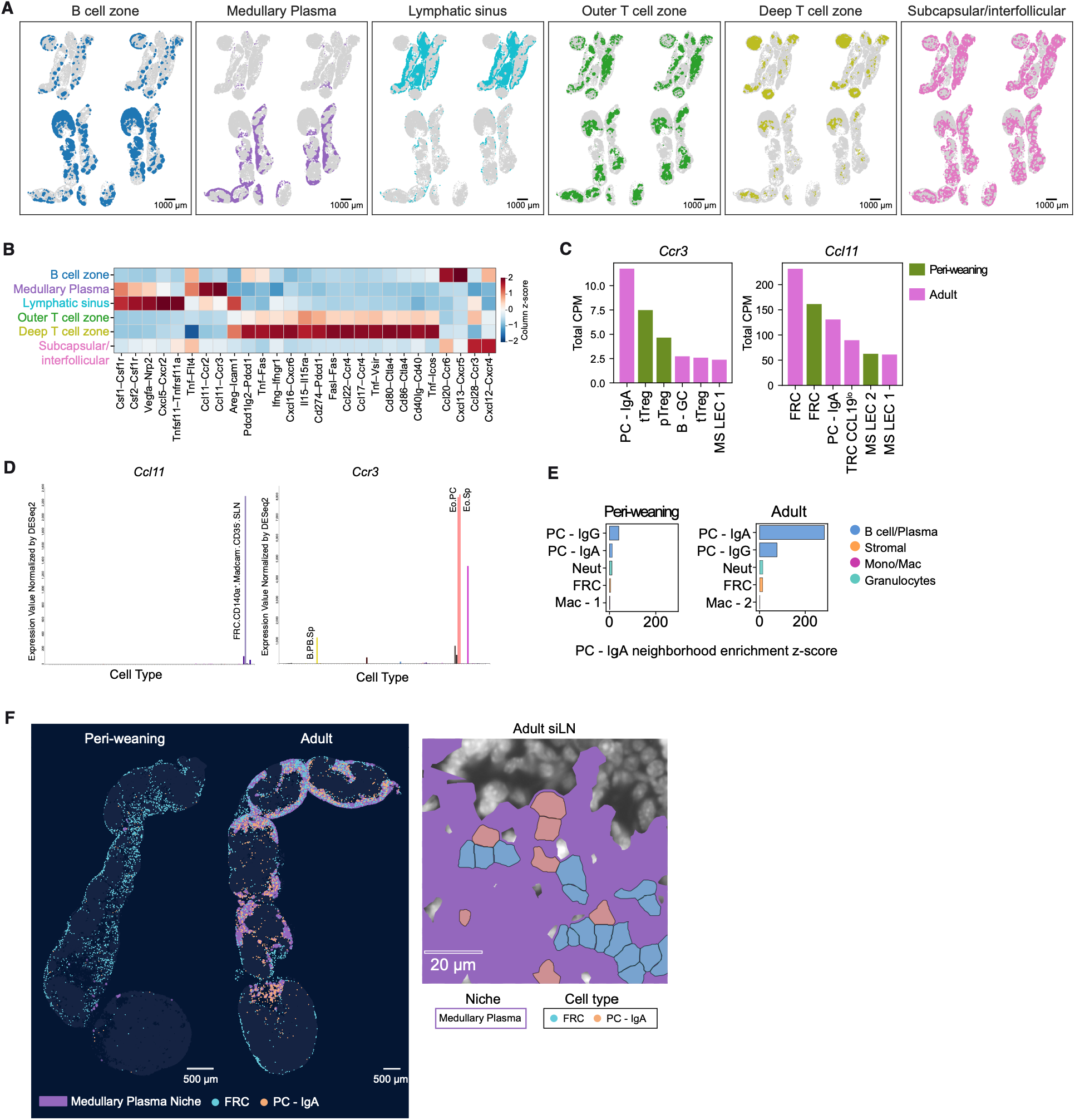
Spatial niches reveal stromal–immune chemokine interactions. (A) Spatial overlays of cellular niches (B cell zone, medullary plasma, subcapsular/interfollicular, lymphatic sinus, deep T cell zone, and outer T cell zone) across all peri-weaning (top) and adult (bottom) mLNs. (B) Heatmap of spatially resolved ligand–receptor activity across niches. (C) Expression of *Ccr3* (left) and *Ccl11* (right) in cell types with the highest expression, shown as total counts per million (CPM), calculated per age group. (D) Expression of *Ccl11* (left) and *Ccr3* (right) across immune and stromal cells profiled by bulk RNA-seq (ImmGen). (E) Neighborhood enrichment of cell types around IgA^+^ plasma cells (PC - IgA) in peri-weaning (top) and adult (bottom) mLN. (F) Spatial distribution of follicular reticular cells (FRCs) and PC - IgA in representative peri-weaning and adult mLNs, colored by cell type and niche (left; scale bars, 500 µm) and a magnified region of an adult siLN (right; scale bar, 20 µm).

**Supplementary Figure 4.**
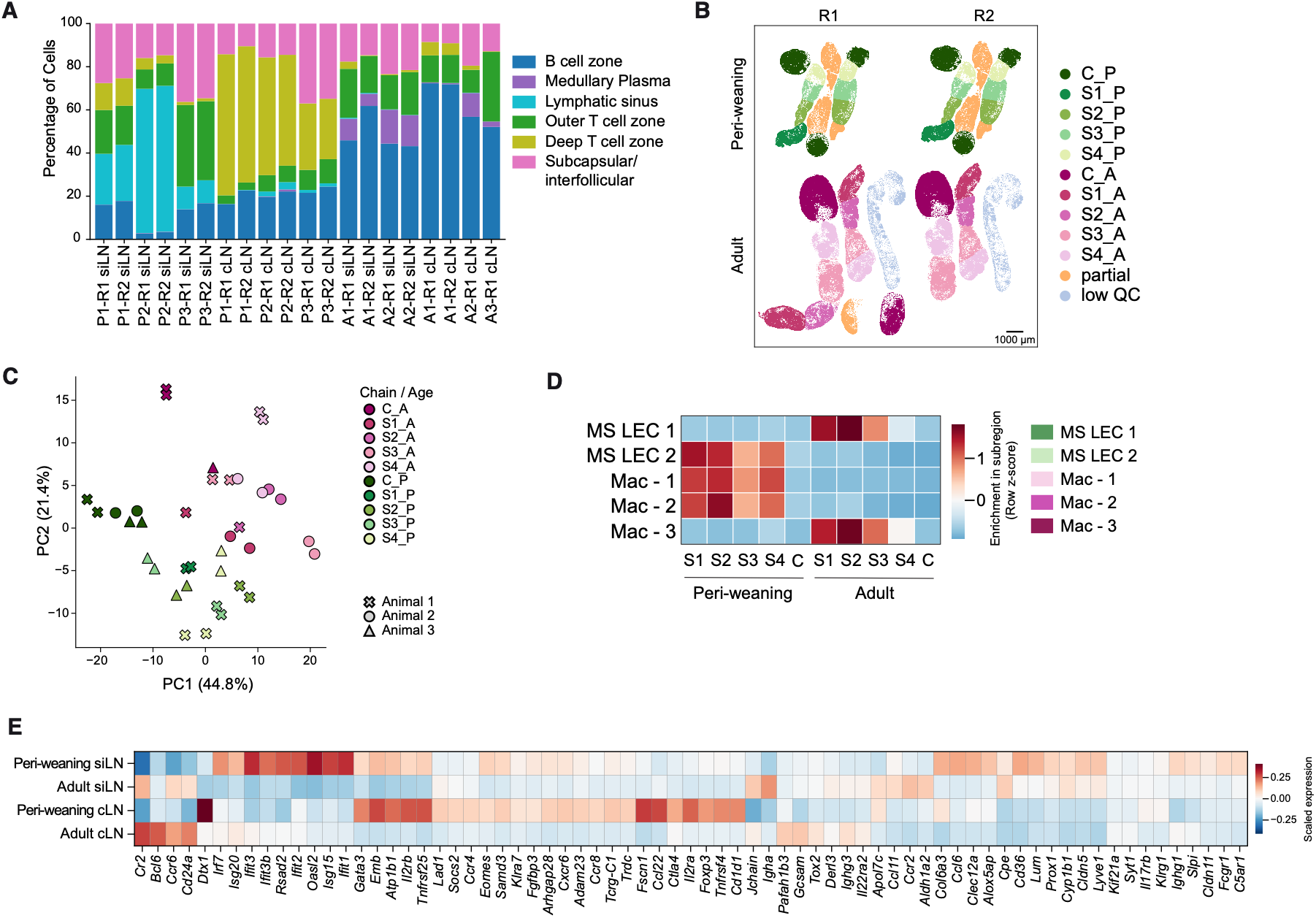
Regional and developmental differences in gut lymph node niche composition. (A) Niche composition for each sample; peri-weaning: P1-P3; adult: A1-A3; Xenium-defined image region: R1, R2; anatomical region: siLN, cLN. (B) mLN chains colored by small intestinal (S1-S4) and colonic (C) draining lymph node subregion. Partial nodes (“partial”) and regions of low-quality tissue (“low QC”) were not included in downstream analysis. (C) Principal component analysis of samples, with each point representing one subregion from one animal and section. (D) Heatmap of the relative abundance of macrophages clusters in peri-weaning and adult mice. (E) Heatmap of DEGs (pseudobulked, PyDESeq2, FDR < 0.05) across age or regional comparisons.

**Supplementary Figure 5.**
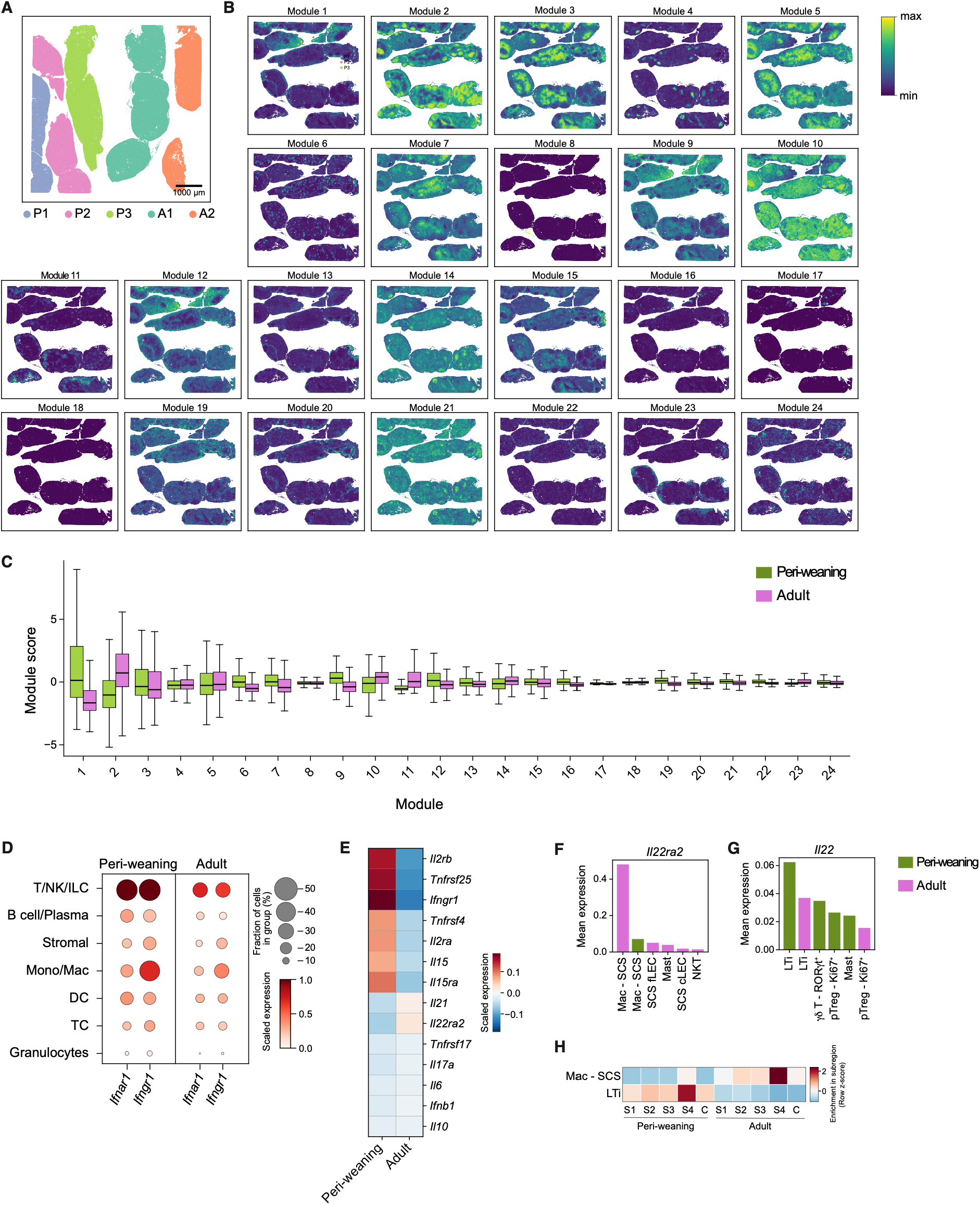
Spatial gene modules and cytokine programs distinguish peri-weaning and adult nodes. (A) mLN sections profiled by Visium HD colored by animal (peri-weaning, P1-P3; adult A1-A2). (B) Spatial overlays of the 24 Hotspot gene module scores. (C) Hotspot module scores in peri-weaning and adult lymph nodes. Box plots show the median and interquartile range (IQR), whiskers extending to 1.5× IQR. (D) *Ifnar1* and *Ifngr1* expression across broad lineages in peri-weaning and adult mice. (E) Heatmap showing scaled expression of differentially expressed cytokine and cytokine receptor genes (PyDESeq2, FDR < 0.05) in peri-weaning and adult lymph nodes. (F, G) Expression of *Il22ra2* (F) and *Il22* (G) in cell types with the highest expression across both adult and peri-weaning mLNs. (H) Heat map of the relative abundance of LTi cells and SSMs (rows) in each mLN subregion.

**Supplementary Figure 6.**
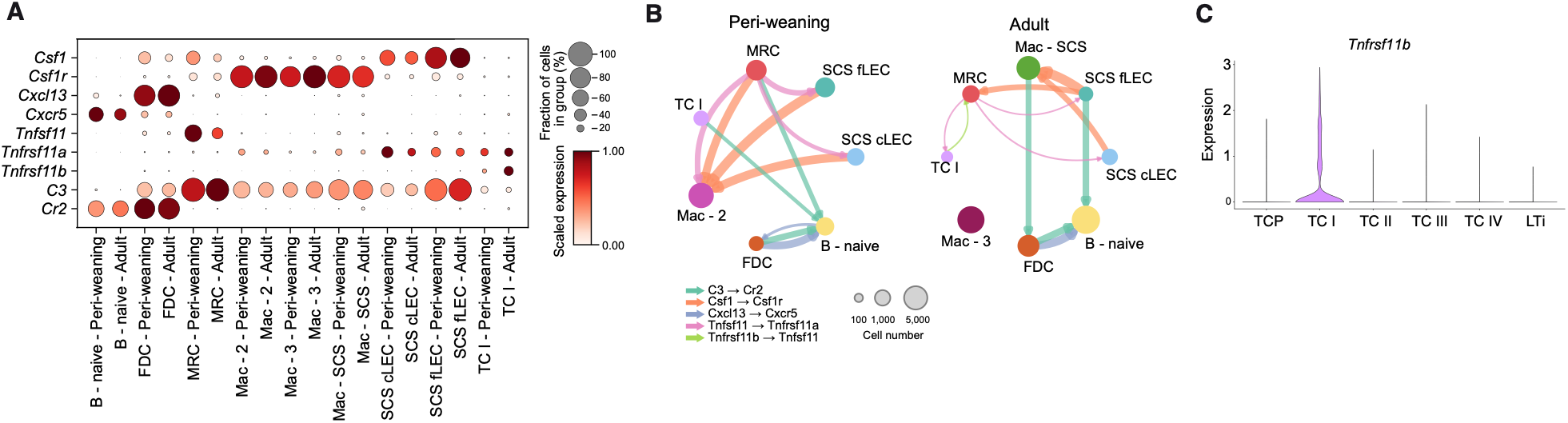
Age-dependent cellular circuits within the subcapsular sinus niche. (A) Dot plot with relative expression of selected ligand (L) and receptor (R) genes for selected cell types in siLN across ages. (B) Cell–cell communication networks for indicated cell types in peri-weaning (left) and adult (right) siLN. Cellphone DB analysis of ligand–receptor interactions colored by L-R pair, with edge width proportional to the mean L-R score. Top 3 interactions (*P* < 0.05) are shown for each pair with self-interactions omitted. (C) Expression of *Tnfrsf11b* in TCs and LTi cells profiled by scRNA-seq (Cabric et al.).

**Supplementary Figure 7.**
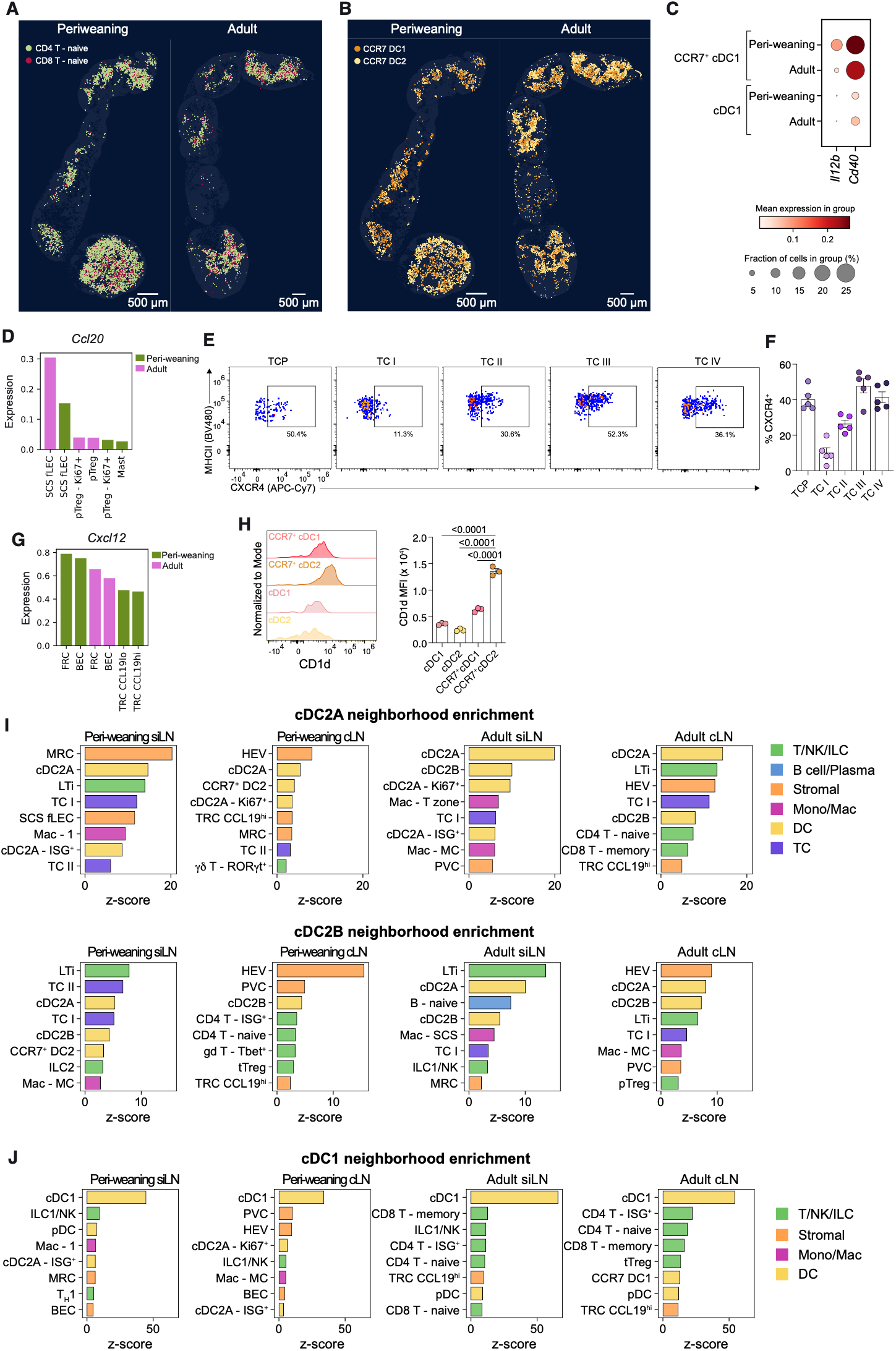
Chemokine expression and cellular neighborhoods of specialized antigen-presenting cells. (A, B) Spatial maps of (A) naive CD4^+^ and CD8^+^ T cells and (B) CCR7^+^ cDC1 and CCR7^+^ cDC2 cells in representative peri-weaning and adult lymph node chains, colored by cell type. (C) Dot plot of *Il12b* and *Cd40* expression in CCR7⁺ cDC1 and cDC1 cells from peri-weaning and adult mice. (D) Expression of *Ccl20* in cell types with the highest expression across adult and peri-weaning mLNs. (E) Representative flow cytometry of CXCR4 expression by TCP and TC subsets in mLNs of P16 *Rorc^Venus-creERT^*^2^ mice and (F) summary graph (n = 5 mice). (G) Expression of *Cxcl12* in the cell types with the highest expression across adult and peri-weaning mLNs. (H) Representative flow cytometry of CD1d expression by cDC subsets in mLNs of P13 *Rorc^Venus^*mice and summary graph (n = 3 mice). Data in E, F, and H are representative of two independent experiments. Error bars: mean ± s.e.m.; one-way ANOVA (H). All *P* values are indicated on the corresponding graphs. (I, J) Neighborhood enrichment of cell types around (I) cDC2A and cDC2B and (J) cDC1 across ages and mLN sub-regions.

**Supplementary Figure 8.**
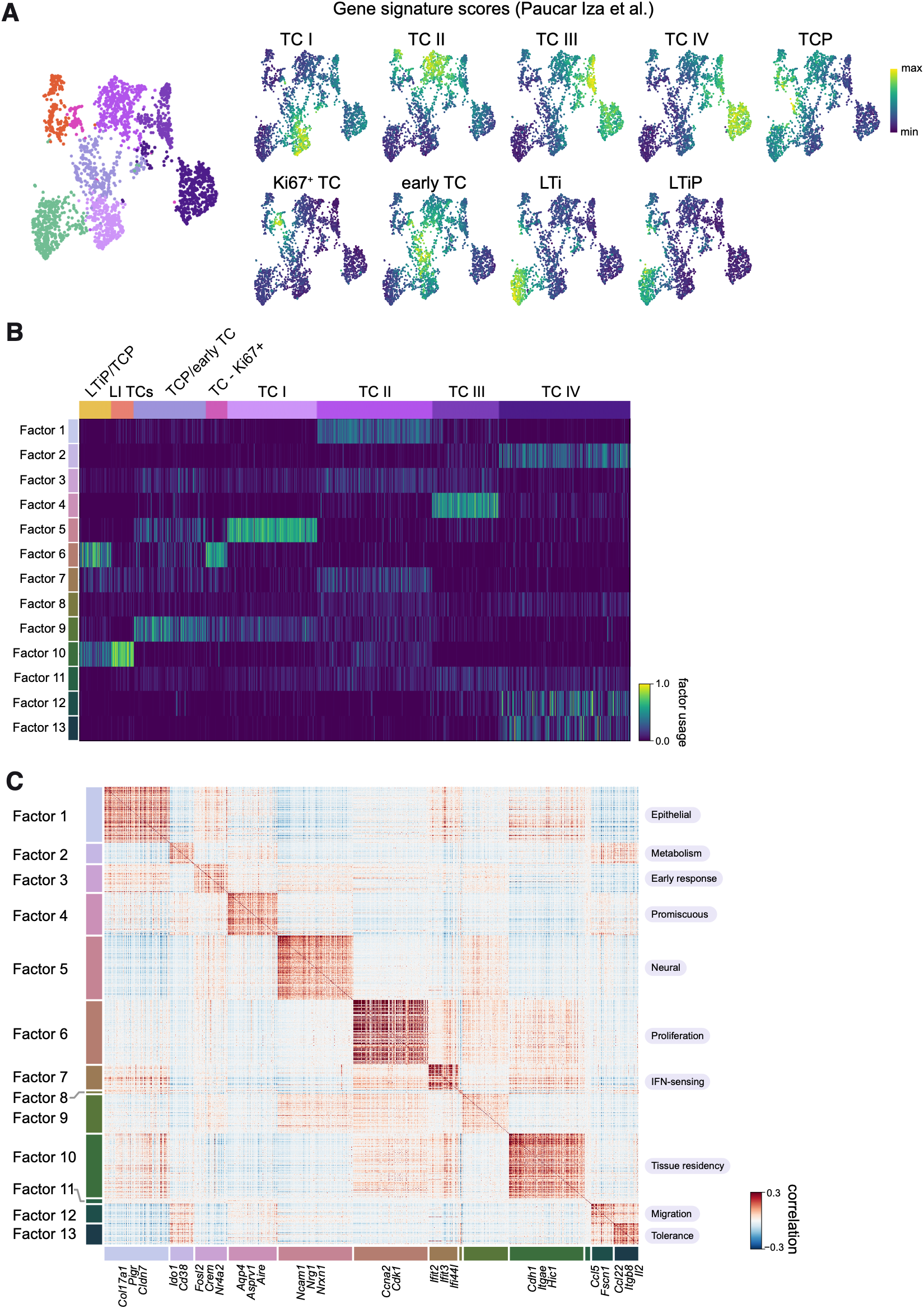
Characterization of intestinal tissue Thetis cells. (A) UMAP of CXCR6^−^RORγt(Venus)^+^ TCs and LTi cells from large intestine and mLNs of P15-18 *Rorc^Venus^* mice profiled by scRNA-seq, colored by cluster or per-cell gene signature scores for indicated cell types (Paucar Iza et al.). (B) Heat map showing relative usage of gene expression programs identified by scHPF (K = 13). (C) Heat map showing scHPF gene scores of the top 200 genes in each ‘factor’ across TCs. Selected genes for each factor are indicated.

**Supplementary Figure 9.**
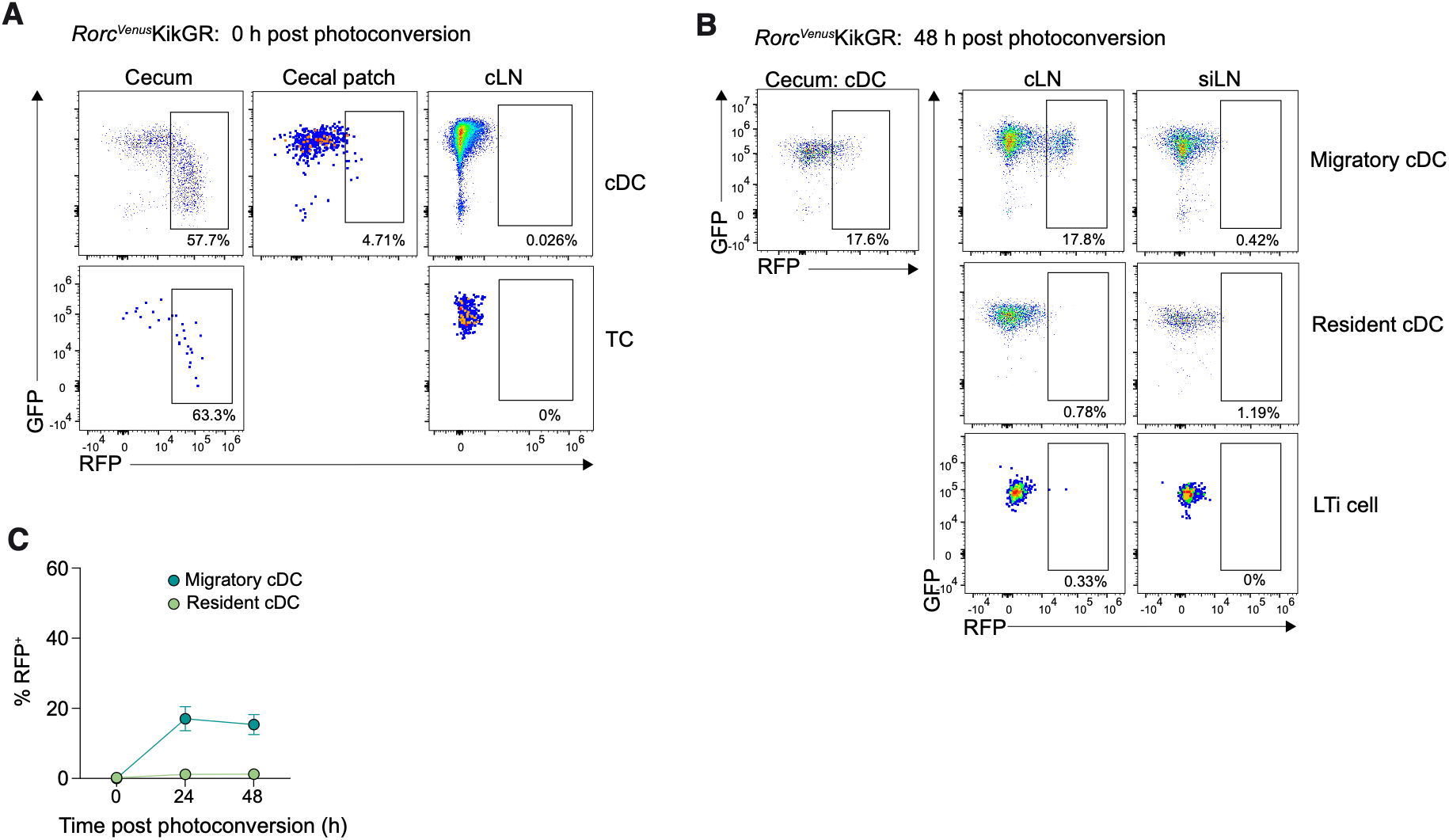
Migration of Thetis cells from tissue to lymph nodes. (A) Representative flow cytometry of RFP expression by cDCs and TCs in cecums, cecal patches and cLNs of P20 *Rorc^Venus^KikGR* mice 0 hr post-photoconversion. (B) Representative flow cytometry of RFP expression by DC subsets and LTi cells in cLNs and siLNs of P20 *Rorc^Venus^KikGR* mice 48 hr post-photoconversion. (C) Summary graph of RFP expression by cDC subsets in cLN of P15-21 *Rorc^Venus^KikGR* mice 48 hr post-photoconversion (0 hr, n = 3; 24 hr, n = 8; 48 hr, n = 7). Data in A, B, and C are pooled from five independent experiments. Error bars: mean ± s.e.m.

## Supplementary Tables

Table S1. Custom Xenium gene panel

Table S2. Flow cytometry antibodies

## Notes

### Competing Interest Statement

The authors have declared no competing interest.

### Summary of Updates

List of authors corrected to reflect manuscript PDF.

## REFERENCES

1. Ohnmacht, C., Park, J.H., Cording, S., Wing, J.B., Atarashi, K., Obata, Y., Gaboriau-Routhiau, V., Marques, R., Dulauroy, S., Fedoseeva, M., et al. (2015). The microbiota regulates type 2 immunity through RORγt+ T cells. Science 349, 989–993. 10.1126/science.aac4263.

2. Xu, M., Pokrovskii, M., Ding, Y., Yi, R., Au, C., Harrison, O.J., Galan, C., Belkaid, Y., Bonneau, R., and Littman, D.R. (2018). C-MAF-dependent regulatory T cells mediate immunological tolerance to a gut pathobiont. Nature 554, 373–377. 10.1038/nature25500.

3. Sefik, E., Geva-Zatorsky, N., Oh, S., Konnikova, L., Zemmour, D., McGuire, A.M., Burzyn, D., Ortiz-Lopez, A., Lobera, M., Yang, J., et al. (2015). Individual intestinal symbionts induce a distinct population of RORγ+ regulatory T cells. Science 349, 993–997. 10.1126/science.aaa9420.

4. Brodin, P. (2022). Immune-microbe interactions early in life: a determinant of health and disease long term. Science 376, 945–950. 10.1126/science.abk2189.

5. Paucar Iza, Y.A., and Brown, C.C. (2024). Early life imprinting of intestinal immune tolerance and tissue homeostasis. Immunol. Rev. 323, 303–315. 10.1111/imr.13321.

6. Du Toit, G., Roberts, G., Sayre, P.H., Bahnson, H.T., Radulovic, S., Santos, A.F., Brough, H.A., Phippard, D., Basting, M., Feeney, M., et al. (2015). Randomized trial of peanut consumption in infants at risk for peanut allergy. N. Engl. J. Med. 372, 803–813. 10.1056/NEJMoa1414850.

7. Jones, S.M., Kim, E.H., Nadeau, K.C., Nowak-Wegrzyn, A., Wood, R.A., Sampson, H.A., Scurlock, A.M., Chinthrajah, S., Wang, J., Pesek, R.D., et al. (2022). Efficacy and safety of oral immunotherapy in children aged 1–3 years with peanut allergy (the immune tolerance network IMPACT trial): a randomised placebo-controlled study. Lancet 399, 359–371. 10.1016/s0140-6736(21)02390-4.

8. Ramanan, D., Sefik, E., Galván-Peña, S., Wu, M., Yang, L., Yang, Z., Kostic, A., Golovkina, T.V., Kasper, D.L., Mathis, D., et al. (2020). An immunologic mode of multigenerational transmission governs a gut Treg setpoint. Cell 181, 1276–1290.e13. 10.1016/j.cell.2020.04.030.

9. Cabric, V., Parisotto, Y.F., Park, T., Akagbosu, B., Zhao, Z., Lo, Y., Shibu, G., Fisher, L., Paucar Iza, Y.A., Leslie, C., et al. (2025). A wave of Thetis cells imparts tolerance to food antigens early in life. Science 389, 268–274. 10.1126/science.adp0535.

10. Kim, K.S., Hong, S.W., Han, D., Yi, J., Jung, J., Yang, B.G., Lee, J.Y., Lee, M., and Surh, C.D. (2016). Dietary antigens limit mucosal immunity by inducing regulatory T cells in the small intestine. Science 351, 858–863. 10.1126/science.aac5560.

11. Nutsch, K., Chai, J.N., Ai, T.L., Russler-Germain, E., Feehley, T., Nagler, C.R., and Hsieh, C.S. (2016). Rapid and efficient generation of regulatory T cells to commensal antigens in the periphery. Cell Rep. 17, 206–220. 10.1016/j.celrep.2016.08.092.

12. Vetter, L., Witchley, J.N., Ahn, S.H., Brewer, R.C., Ly, A., Tout, A., Wagner, A., and Barton, G.M. (2026). Profiling of intestinal T cells reveals antigen-specific dependency on early-life programming for the generation of microbiota-reactive Tregs. bioRxiv 10.64898/2026.09.23.753882.

13. Chang, Y.-T., Le, H.H., Morrison, A., Kong, R., Schulman, E., Chou, T.-W., and Blum, J.E. (2026). Intestinal uptake regulates T cell responses to dietary antigens. bioRxiv 10.64898/2026.09.01.748748.

14. Skjerven, H.O., Lie, A., Vettukattil, R., Rehbinder, E.M., LeBlanc, M., Asarnoj, A., Carlsen, K.H., Despriee, Å.W., Färdig, M., Gerdin, S.W., et al. (2022). Early food intervention and skin emollients to prevent food allergy in young children (PreventADALL): a factorial, multicentre, cluster-randomised trial. Lancet 399, 2398–2411. 10.1016/s0140-6736(22)00687-0.

15. Akagbosu, B., Tayyebi, Z., Shibu, G., Paucar Iza, Y.A., Deep, D., Parisotto, Y.F., Fisher, L., Pasolli, H.A., Thevin, V., Elmentaite, R., et al. (2022). Novel antigen-presenting cell imparts treg-dependent tolerance to gut microbiota. Nature 610, 752–760. 10.1038/s41586-022-05309-5.

16. Paucar Iza, Y.A., Park, T., Baker, E., Shibu, G., Hoelting, T., Feather, G., Yadav, A., Parisotto, Y.F., Zhao, Z., Akagbosu, B., et al. (2026). Ontogeny and transcriptional regulation of Thetis cells. Nature 653, 538–547. 10.1038/s41586-026-10198-z.

17. Rodrigues, P.F., Wu, S., Trsan, T., Bhattarai, B., Panda, S.K., Fachi, J.L., Cella, M., and Colonna, M. (2026). RORγt+ dendritic cells are a distinct lymphoid-derived lineage. Sci. Immunol. 11, eaed7439. 10.1126/sciimmunol.aed7439.

18. Rodrigues, P.F., Wu, S., Trsan, T., Panda, S.K., Fachi, J.L., Liu, Y., Du, S., de Oliveira, S., Ulezko Antonova, A., Khantakova, D., et al. (2025). Rorγt-positive dendritic cells are required for the induction of peripheral regulatory T cells in response to oral antigens. Cell 188, 2720–2737.e22. 10.1016/j.cell.2025.03.020.

19. Kedmi, R., Najar, T.A., Mesa, K.R., Grayson, A., Kroehling, L., Hao, Y., Hao, S., Pokrovskii, M., Xu, M., Talbot, J., et al. (2022). A RORγt+ cell instructs gut microbiota-specific Treg cell differentiation. Nature 610, 737–743. 10.1038/s41586-022-05089-y.

20. Fu, L., Upadhyay, R., Pokrovskii, M., Chen, F.M., Romero-Meza, G., Griesemer, A., and Littman, D.R. (2025). PRDM16-dependent antigen-presenting cells induce tolerance to gut antigens. Nature 642, 756–765. 10.1038/s41586-025-08982-4.

21. Rudnitsky, A., Oh, H., Margolin, M., Dassa, B., Shteinberg, I., Stoler-Barak, L., Shulman, Z., and Kedmi, R. (2025). A coordinated cellular network regulates tolerance to food. Nature 644, 231–240. 10.1038/s41586-025-09173-x.

22. Sun, I.H., Qualls, A.E., Yin, H.S., Wang, J., Arvedson, M.P., Germino, J., Horner, N.K., Zhong, S., Du, J., Valdearcos, M., et al. (2025). RORγt eTACs mediate oral tolerance and Treg induction. J. Exp. Med. 222, e20250573. 10.1084/jem.20250573.

23. Chi, X., Wang, C.H., Parisotto, Y.F., Nyberg, W.A., Cabric, V., Gelineau, A., Cao, Y., Owen, D.L., Ambjörnsson, J., Mathis, D., et al. (2026). TCR origin and specificity to environmental or self-antigens on distinct antigen-presenting cells determine peripheral Treg cell differentiation. Immunity 59, 2266–2281.e6. 10.1016/j.immuni.2026.07.001.

24. Zhou, Y.D., Brown, H., Schaffer, E., Taylor, G.M., Fiske, K.L., Komnick, M.R., Lopez, S., Dermody, T.S., and Esterházy, D. (2026). T cell fate is dictated by different antigen-presenting cells in response to dietary versus gut epithelial self-antigen. Immunity 59, 2758–2774.e9. 10.1016/j.immuni.2026.07.023.

25. Blum, J.E., Kong, R., Schulman, E., Chen, F.M., Upadhyay, R., Romero-Meza, G., Littman, D.R., Fischbach, M.A., Nagashima, K., and Sattely, E.S. (2026). Identification and characterization of dietary antigens in oral tolerance. Sci. Immunol. 11, eaeb4684. 10.1126/sciimmunol.aeb4684.

26. Hoelting, T.L., Park, T., and Brown, C.C. (2025). Antigen-presenting cells as arbiters of mucosal tolerance and immunity. Nat. Immunol. 26, 1890–1902. 10.1038/s41590-025-02320-6.

27. Narasimhan, H., Richter, M.L., Shakiba, R., Papaioannou, N.E., Stehle, C., Ravi Rengarajan, K., Ulmert, I., Kendirli, A., de la Rosa, C., Kuo, P.Y., et al. (2025). RORγt-expressing dendritic cells are functionally versatile and evolutionarily conserved antigen-presenting cells. Proc. Natl. Acad. Sci. USA 122, e2417308122. 10.1073/pnas.2417308122.

28. Torow, N., Li, R., Charles, T., Hitch, A., Lelouard, H., Gesteira Costa, I., and Hornef, M.W. (2023). M cell maturation and cDC activation determine the onset of adaptive immune priming in the neonatal Peyer’s patch. Immunity 56, 1220–1238.e7. 10.1016/j.immuni.2023.04.002.

29. Roozendaal, R., Mempel, T.R., Pitcher, L.A., Gonzalez, S.F., Verschoor, A., Mebius, R.E., von Andrian, U.H., and Carroll, M.C. (2009). Conduits mediate transport of low-molecular-weight antigen to lymph node follicles. Immunity 30, 264–276. 10.1016/j.immuni.2008.12.014.

30. Goenka, R., Barnett, L.G., Silver, J.S., O’Neill, P.J., Hunter, C.A., Cancro, M.P., and Laufer, T.M. (2011). Cutting edge: dendritic Cell-Restricted antigen presentation initiates the follicular helper T cell program but cannot complete ultimate effector differentiation. J. Immunol. 187, 1091–1095. 10.4049/jimmunol.1100853.

31. Barnett, L.G., Simkins, H.M.A., Barnett, B.E., Korn, L.L., Johnson, A.L., Wherry, E.J., Wu, G.F., and Laufer, T.M. (2014). B cell antigen presentation in the initiation of follicular helper T cell and germinal center differentiation. J. Immunol. 192, 3607–3617. 10.4049/jimmunol.1301284.

32. Phan, T.G., Grigorova, I., Okada, T., and Cyster, J.G. (2007). Subcapsular encounter and complement-dependent transport of immune complexes by lymph node B cells. Nat. Immunol. 8, 992–1000. 10.1038/ni1494.

33. Phan, T.G., Green, J.A., Xu, Y., and Cyster, J.G. (2009). Immune complex relay by subcapsular sinus macrophages and noncognate B cells drives antibody affinity maturation. Nat. Immunol. 10, 786–796. 10.1038/ni.1745.

34. Gerner, M.Y., Casey, K.A., Kastenmuller, W., and Germain, R.N. (2017). Dendritic cell and antigen dispersal landscapes regulate T cell immunity. J. Exp. Med. 214, 3105–3122. 10.1084/jem.20170335.

35. Sixt, M., Kanazawa, N., Selg, M., Samson, T., Roos, G., Reinhardt, D.P., Pabst, R., Lutz, M.B., and Sorokin, L. (2005). The conduit system transports soluble antigens from the afferent lymph to resident dendritic cells in the T cell area of the lymph node. Immunity 22, 19–29. 10.1016/j.immuni.2004.11.013.

36. Gerner, M.Y., Torabi-Parizi, P., and Germain, R.N. (2015). Strategically localized dendritic cells promote rapid T cell responses to lymph-borne particulate antigens. Immunity 42, 172–185. 10.1016/j.immuni.2014.12.024.

37. Martín-Fontecha, A., Sebastiani, S., Höpken, U.E., Uguccioni, M., Lipp, M., Lanzavecchia, A., and Sallusto, F. (2003). Regulation of dendritic cell migration to the draining lymph node: impact on T lymphocyte traffic and priming. J. Exp. Med. 198, 615–621. 10.1084/jem.20030448.

38. Förster, R., Schubel, A., Breitfeld, D., Kremmer, E., Renner-Müller, I., Wolf, E., and Lipp, M. (1999). CCR7 coordinates the primary immune response by establishing functional microenvironments in secondary lymphoid organs. Cell 99, 23–33. 10.1016/s0092-8674(00)80059-8.

39. Pezoldt, J., Pasztoi, M., Zou, M., Wiechers, C., Beckstette, M., Thierry, G.R., Vafadarnejad, E., Floess, S., Arampatzi, P., Buettner, M., et al. (2018). Neonatally imprinted stromal cell subsets induce tolerogenic dendritic cells in mesenteric lymph nodes. Nat. Commun. 9, 3903. 10.1038/s41467-018-06423-7.

40. Heidari, E., Moorman, A., Unyi, D., Pasnuri, N., Rukhovich, G., Calafato, D., Mathioudaki, A., Chan, J.M., Nawy, T., Gerstung, M., et al. (2025). Segger: fast and accurate cell segmentation of imaging-based spatial transcriptomics data. bioRxiv 10.1101/2025.03.14.643160.

41. Huang, H.Y., Rivas-Caicedo, A., Renevey, F., Cannelle, H., Peranzoni, E., Scarpellino, L., Hardie, D.L., Pommier, A., Schaeuble, K., Favre, S., et al. (2018). Identification of a new subset of lymph node stromal cells involved in regulating plasma cell homeostasis. Proc. Natl. Acad. Sci. USA 115, E6826–E6835. 10.1073/pnas.1712628115.

42. Esterházy, D., Canesso, M.C., Mesin, L., Muller, P.A., de Castro, T.B., Lockhart, A., ElJalby, M., Faria, A.M., and Mucida, D. (2019). Compartmentalized gut lymph node drainage dictates adaptive immune responses. Nature 569, 126–130. 10.1038/s41586-019-1125-3.

43. Fatkhullina, A.R., Kent, J., Brown, H., Christiansen, N., Lisicka, W., Madariaga, M.L., and Esterházy, D. (2025). Medullary stromal cells define small intestinal lymph node identity in humans and mice. Cell Rep. 44, 116441. 10.1016/j.celrep.2025.116441.

44. Rusconi, B., Bard, A.K., McDonough, R., Kindsvogel, A.M., Wang, J.D., Udayan, S., McDonald, K.G., Newberry, R.D., and Tarr, P.I. (2024). Intergenerational protective anti-gut commensal immunoglobulin g originates in early life. Proc. Natl. Acad. Sci. USA 121, e2309994121. 10.1073/pnas.2309994121.

45. Bos, A., Egmond, M., and Mebius, R. (2022). The role of retinoic acid in the production of immunoglobulin A. Mucosal Immunol. 15, 562–572. 10.1038/s41385-022-00509-8.

46. DeTomaso, D., and Yosef, N. (2021). Hotspot identifies informative gene modules across modalities of single-cell genomics. Cell Syst. 12, 446–456.e9. 10.1016/j.cels.2021.04.005.

47. Al Nabhani, Z., Dulauroy, S., Marques, R., Cousu, C., Al Bounny, S., Déjardin, F., Sparwasser, T., Bérard, M., Cerf-Bensussan, N., and Eberl, G. (2019). A weaning reaction to microbiota is required for resistance to immunopathologies in the adult. Immunity 50, 1276–1288.e5. 10.1016/j.immuni.2019.02.014.

48. Hart, A., Merolle, M., Howard, C., Haskins, B.E., Cohn, I.S., Bobba, S., Xiao, R., Yang, Y., Cadwell, K., Ma, J., et al. (2026). Diverse infections transcriptionally reprogram the intestinal epithelium and epithelial–immune cell interactions. Nat. Immunol. 10.1038/s41590-026-02665-6.

49. Jinnohara, T., Kanaya, T., Hase, K., Sakakibara, S., Kato, T., Tachibana, N., Sasaki, T., Hashimoto, Y., Sato, T., Watarai, H., et al. (2017). IL-22BP dictates characteristics of Peyer’s patch follicleassociated epithelium for antigen uptake. J. Exp. Med. 214, 1607–1618. 10.1084/jem.20160770.

50. Barone, F., Nayar, S., Campos, J., Cloake, T., Withers, D.R., Toellner, K.-M., Zhang, Y., Fouser, L., Fisher, B., Bowman, S., et al. (2015). IL-22 regulates lymphoid chemokine production and assembly of tertiary lymphoid organs. Proc. Natl. Acad. Sci. USA 112, 11024–11029. 10.1073/pnas.1503315112.

51. Udayan, S., Floyd, A.N., John, V., Barrios, B.E., Rusconi, B.A., McDonald, K.G., Schill, E.M., Kulkarni, D.H., Martin, A.L., Gutierrez, R., et al. (2025). Colonic goblet cell-associated antigen passages mediate physiologic and beneficial translocation of live gut bacteria in preweaning mice. Nat. Microbiol. 10, 927–938. 10.1038/s41564-025-01965-1.

52. Harris, N.L., Spoerri, I., Schopfer, J.F., Nembrini, C., Merky, P., Massacand, J., Urban, J.F., Lamarre, A., Burki, K., Odermatt, B., et al. (2006). Mechanisms of neonatal mucosal antibody protection. J. Immunol. 177, 6256–6262. 10.4049/jimmunol.177.9.6256.

53. Mebius, R.E., and Kraal, G. (2005). Structure and function of the spleen. Nat. Rev. Immunol. 5, 606–616. 10.1038/nri1669.

54. Mondor, I., Baratin, M., Lagueyrie, M., Saro, L., Henri, S., Gentek, R., Suerinck, D., Kastenmuller, W., Jiang, J.X., and Bajénoff, M. (2019). Lymphatic endothelial cells are essential components of the subcapsular sinus macrophage niche. Immunity 50, 1453–1466.e4. 10.1016/j.immuni.2019.04.002.

55. Kang, Y.S., Do, Y., Lee, H.K., Park, S.H., Cheong, C., Lynch, R.M., Loeffler, J.M., Steinman, R.M., and Park, C.G. (2006). A dominant complement fixation pathway for pneumococcal polysaccharides initiated by SIGN-R1 interacting with C1q. Cell 125, 47–58. 10.1016/j.cell.2006.01.046.

56. Camara, A., Cordeiro, O.G., Alloush, F., Sponsel, J., Chypre, M., Onder, L., Asano, K., Tanaka, M., Yagita, H., Ludewig, B., et al. (2019). Lymph node mesenchymal and endothelial stromal cells cooperate via the RANK-RANKL cytokine axis to shape the sinusoidal macrophage niche. Immunity 50, 1467–1481.e6. 10.1016/j.immuni.2019.05.008.

57. Camara, A., Lavanant, A.C., Abe, J., Desforges, H.L., Alexandre, Y.O., Girardi, E., Igamberdieva, Z., Asano, K., Tanaka, M., Hehlgans, T., et al. (2022). CD169+ macrophages in lymph node and spleen critically depend on dual RANK and LTbetaR signaling. Proc. Natl. Acad. Sci. USA 119, e2108540119. 10.1073/pnas.2108540119.

58. Moseman, E.A., Iannacone, M., Bosurgi, L., Tonti, E., Chevrier, N., Tumanov, A., Fu, Y.X., Hacohen, N., and von Andrian, U.H. (2012). B cell maintenance of subcapsular sinus macrophages protects against a fatal viral infection independent of adaptive immunity. Immunity 36, 415–426. 10.1016/j.immuni.2012.01.013.

59. Koch, M.A., Reiner, G.L., Lugo, K.A., Kreuk, L.S., Stanbery, A.G., Ansaldo, E., Seher, T.D., Ludington, W.B., and Barton, G.M. (2016). Maternal IgG and IgA antibodies dampen mucosal T helper cell responses in early life. Cell 165, 827–841. 10.1016/j.cell.2016.04.055.

60. Alouche, N., Cannelle, H., Favre, S., Jokić, M., Villegas, J.A., Maréchal, S., De Nolf, C., Delacrétaz, M., Sifis, A., Chapuis, C., et al. (2026). Homeostatic mature dendritic cells instruct fibroblast specialization via Notch2 signaling to establish T cell niches. Immunity 59, 1688–1707.e11. 10.1016/j.immuni.2026.03.023.

61. Lu, E., Dang, E.V., McDonald, J.G., and Cyster, J.G. (2017). Distinct oxysterol requirements for positioning naïve and activated dendritic cells in the spleen. Sci. Immunol. 2, eaal5237. 10.1126/sciimmunol.aal5237.

62. Pihlgren, M., Tougne, C., Bozzotti, P., Fulurija, A., Duchosal, M.A., Lambert, P.-H., and Siegrist, C.-A. (2003). Unresponsiveness to Lymphoid-Mediated signals at the neonatal follicular dendritic cell precursor level contributes to delayed germinal center induction and limitations of neonatal antibody responses to T-Dependent antigens. J. Immunol. 170, 2824–2832. 10.4049/jimmunol.170.6.2824.

63. Mastelic, B., Kamath, A.T., Fontannaz, P., Tougne, C., Rochat, A.-F., Belnoue, E., Combescure, C., Auderset, F., Lambert, P.-H., Tacchini-Cottier, F., et al. (2012). Environmental and T cell–intrinsic factors limit the expansion of neonatal follicular T helper cells but may be circumvented by specific adjuvants. J. Immunol. 189, 5764–5772. 10.4049/jimmunol.1201143.

64. Debock, I., Jaworski, K., Chadlaoui, H., Delbauve, S., Passon, N., Twyffels, L., Leo, O., and Flamand, V. (2013). Neonatal follicular Th cell responses are impaired and modulated by IL-4. J. Immunol. 191, 1231–1239. 10.4049/jimmunol.1203288.

65. Kramer, D.R., and Cebra, J.J. (1995). Early appearance of “natural” mucosal IgA responses and germinal centers in suckling mice developing in the absence of maternal antibodies. J. Immunol. 154, 2051–2062. https://pubmed.ncbi.nlm.nih.gov/7868882/.

66. Neumann, C., Blume, J., Roy, U., Teh, P.P., Vasanthakumar, A., Beller, A., Liao, Y., Heinrich, F., Arenzana, T.L., Hackney, J.A., et al. (2019). c-Maf-dependent Treg cell control of intestinal TH17 cells and IgA establishes host–microbiota homeostasis. Nat. Immunol. 20, 471–481. 10.1038/s41590-019-0316-2.

67. Knoop, K.A., McDonald, K.G., McCrate, S., McDole, J.R., and Newberry, R.D. (2014). Microbial sensing by goblet cells controls immune surveillance of luminal antigens in the colon. Mucosal Immunol. 8, 198–210. 10.1038/mi.2014.58.

68. Knoop, K.A., Gustafsson, J.K., McDonald, K.G., Kulkarni, D.H., Coughlin, P.E., McCrate, S., Kim, D., Hsieh, C.S., Hogan, S.P., Elson, C.O., et al. (2017). Microbial antigen encounter during a preweaning interval is critical for tolerance to gut bacteria. Sci. Immunol. 2, eaao1314. 10.1126/sciimmunol.aao1314.

69. Ulezko Antonova, A., Lonardi, S., Monti, M., Missale, F., Fan, C., Coates, M.L., Bugatti, M., Jaeger, N., Fernandes Rodrigues, P., Brioschi, S., et al. (2023). A distinct human cell type expressing MHCII and RORγt with dual characteristics of dendritic cells and type 3 innate lymphoid cells. Proc. Natl. Acad. Sci. USA 120, e2318710120. 10.1073/pnas.2318710120.

70. Blum, S.M., Chan, T.L., Smith, N.P., Ambrose, C., Xu, K.H., Tirard, A., Samanta, N., Martin, S., Best, R., Tuttle, E., et al. (2026). Multiomic analysis of malignant ascites defines RORC-expressing dendritic cells at sites of metastasis. Sci. Immunol. 11, eaeb9068. 10.1126/sciimmunol.aeb9068.

71. Griswold, S.L., Sajja, K.C., Jang, C.W., and Behringer, R.R. (2011). Generation and characterization of iUBC-KikGR photoconvertible transgenic mice for live time-lapse imaging during development. Genesis 49, 591–598. 10.1002/dvg.20718.

72. Marconato, L., Palla, G., Yamauchi, K.A., Virshup, I., Heidari, E., Treis, T., Vierdag, W.M., Toth, M., Stockhaus, S., Shrestha, R.B., et al. (2025). SpatialData: an open and universal data framework for spatial omics. Nat. Methods 22, 58–62. 10.1038/s41592-024-02212-x.

73. Wolf, F.A., Angerer, P., and Theis, F.J. (2018). SCANPY: large-scale single-cell gene expression data analysis. Genome Biol. 19, 15. 10.1186/s13059-017-1382-0.

74. Levine, J.H., Simonds, E.F., Bendall, S.C., Davis, K.L., Amir, E.-A.D., Tadmor, M.D., Litvin, O., Fienberg, H.G., Jager, A., Zunder, E.R., et al. (2015). Data-Driven phenotypic dissection of AML reveals progenitor-like cells that correlate with prognosis. Cell 162, 184–197. 10.1016/j.cell.2015.05.047.

75. Brown, C.C., Gudjonson, H., Pritykin, Y., Deep, D., Lavallée, V.P., Mendoza, A., Fromme, R., Mazutis, L., Ariyan, C., Leslie, C., et al. (2019). Transcriptional basis of mouse and human dendritic cell heterogeneity. Cell 179, 846–863.e24. 10.1016/j.cell.2019.09.035.

76. Rodrigues, P.F., Trsan, T., Cvijetic, G., Khantakova, D., Panda, S.K., Liu, Z., Ginhoux, F., Cella, M., and Colonna, M. (2024). Progenitors of distinct lineages shape the diversity of mature type 2 conventional dendritic cells. Immunity 57, 1567–1585.e5. 10.1016/j.immuni.2024.05.007.

77. Malhotra, D., Fletcher, A.L., Astarita, J., Lukacs-Kornek, V., Tayalia, P., Gonzalez, S.F., Elpek, K.G., Chang, S.K., Knoblich, K., Hemler, M.E., et al. (2012). Transcriptional profiling of stroma from inflamed and resting lymph nodes defines immunological hallmarks. Nat. Immunol. 13, 499–510. 10.1038/ni.2262.

78. Rodda, L.B., Lu, E., Bennett, M.L., Sokol, C.L., Wang, X., Luther, S.A., Barres, B.A., Luster, A.D., Ye, C.J., and Cyster, J.G. (2018). Single-Cell RNA sequencing of lymph node stromal cells reveals Niche-Associated heterogeneity. Immunity 48, 1014–1028.e6. 10.1016/j.immuni.2018.04.006.

79. Xiang, M., Grosso, R.A., Takeda, A., Pan, J., Bekkhus, T., Brulois, K., Dermadi, D., Nordling, S., Vanlandewijck, M., Jalkanen, S., et al. (2020). A Single-Cell transcriptional roadmap of the mouse and human lymph node lymphatic vasculature. Front. Cardiovasc. Med. 7, 52. 10.3389/fcvm.2020.00052.

80. Nguyen, A.A., Fisher, L., Ahn, J.S., Zuo, M., Park, T., Yadav, A., Gams, M.S., Haniuda, K., Dasoveanu, D.C., Li, C., et al. (2025). Sequential lymphotoxin-β receptor and retinoic acid receptor signals regulate cDC2 fate. Nat. Immunol. 26, 2159–2169. 10.1038/s41590-025-02329-x.

81. van der Veeken, J., Campbell, C., Pritykin, Y., Schizas, M., Verter, J., Hu, W., Wang, Z.M., Matheis, F., Mucida, D., Charbonnier, L.M., et al. (2022). Genetic tracing reveals transcription factor Foxp3-dependent and Foxp3-independent functionality of peripherally induced Treg cells. Immunity 55, 1173–1184.e7. 10.1016/j.immuni.2022.05.010.

82. Ramanan, D., Chowdhary, K., Candéias, S.M., Sassone-Corsi, M., Gelineau, A., Mathis, D., and Benoist, C. (2023). Homeostatic, repertoire and transcriptional relationships between colon T regulatory cell subsets. Proc. Natl. Acad. Sci. USA 120, e2311566120. 10.1073/pnas.2311566120.

83. Ramanan, D., Pratama, A., Zhu, Y., Venezia, O., Sassone-Corsi, M., Chowdhary, K., Galván-Peña, S., Sefik, E., Brown, C., Gélineau, A., et al. (2023). Regulatory T cells in the face of the intestinal microbiota. Nat. Rev. Immunol. 23, 749–762. 10.1038/s41577-023-00890-w.

84. Grueter, B., Petter, M., Egawa, T., Laule-Kilian, K., Aldrian, C.J., Wuerch, A., Ludwig, Y., Fukuyama, H., Wardemann, H., Waldschuetz, R., et al. (2005). Runx3 regulates integrin αe/CD103 and CD4 expression during development of CD4−/CD8+ T cells. J. Immunol. 175, 1694–1705. 10.4049/jimmunol.175.3.1694.

85. Istaces, N., Splittgerber, M., Lima Silva, V., Nguyen, M., Thomas, S., Le, A., Achouri, Y., Calonne, E., Defrance, M., Fuks, F., et al. (2019). EOMES interacts with RUNX3 and BRG1 to promote innate memory cell formation through epigenetic reprogramming. Nat. Commun. 10, 3306. 10.1038/s41467-019-11233-6.

86. Coles, M.C., McMahon, C.W., Takizawa, H., and Raulet, D.H. (2000). Memory CD8 T lymphocytes express inhibitory MHC-specific Ly49 receptors. Eur. J. Immunol. 30, 236–244. 10.1002/1521-4141(200001)30:1<236::AID-IMMU236>3.0.CO;2-X.

87. Baratin, M., Simon, L., Jorquera, A., Ghigo, C., Dembele, D., Nowak, J., Gentek, R., Wienert, S., Klauschen, F., Malissen, B., et al. (2017). T cell zone resident macrophages silently dispose of apoptotic cells in the lymph node. Immunity 47, 349–362.e5. 10.1016/j.immuni.2017.07.019.

88. Phipson, B., Sim, C.B., Porrello, E.R., Hewitt, A.W., Powell, J., and Oshlack, A. (2022). propeller: testing for differences in cell type proportions in single cell data. Bioinformatics 38, 4720–4726. 10.1093/bioinformatics/btac582.

89. Love, M.I., Huber, W., and Anders, S. (2014). Moderated estimation of fold change and dispersion for RNA-seq data with DESeq2. Genome Biol. 15, 550. 10.1186/s13059-014-0550-8.

90. Badia-I-Mompel, P., Vélez Santiago, J., Braunger, J., Geiss, C., Dimitrov, D., Müller-Dott, S., Taus, P., Dugourd, A., Holland, C.H., Ramirez Flores, R.O., et al. (2022). decoupleR: ensemble of computational methods to infer biological activities from omics data. Bioinform. Adv. 2, vbac016. 10.1093/bioadv/vbac016.

91. Dimitrov, D., Schäfer, P.S.L., Farr, E., Rodriguez-Mier, P., Lobentanzer, S., Badia-i-Mompel, P., Dugourd, A., Tanevski, J., Ramirez Flores, R.O., and Saez-Rodriguez, J. (2024). LIANA+ provides an all-in-one framework for cell–cell communication inference. Nat. Cell Biol. 26, 1613–1622. 10.1038/s41556-024-01469-w.

92. Efremova, M., Vento-Tormo, M., Teichmann, S.A., and Vento-Tormo, R. (2020). CellPhoneDB: inferring cell–cell communication from combined expression of multi-subunit ligand–receptor complexes. Nat. Protoc. 15, 1484–1506. 10.1038/s41596-020-0292-x.

93. Palla, G., Spitzer, H., Klein, M., Fischer, D., Schaar, A.C., Kuemmerle, L.B., Rybakov, S., Ibarra, I.L., Holmberg, O., Virshup, I., et al. (2022). Squidpy: a scalable framework for spatial omics analysis. Nat. Methods 19, 171–178. 10.1038/s41592-021-01358-2.

94. Setty, M., Kiseliovas, V., Levine, J., Gayoso, A., Mazutis, L., and Pe’er, D. (2019). Characterization of cell fate probabilities in single-cell data with palantir. Nat. Biotechnol. 37, 451–460. 10.1038/s41587-019-0068-4.

95. Finak, G., McDavid, A., Yajima, M., Deng, J., Gersuk, V., Shalek, A.K., Slichter, C.K., Miller, H.W., McElrath, M.J., Prlic, M., et al. (2015). MAST: a flexible statistical framework for assessing transcriptional changes and characterizing heterogeneity in single-cell RNA sequencing data. Genome Biol. 16, 278. 10.1186/s13059-015-0844-5.

96. van Dijk, D., Sharma, R., Nainys, J., Yim, K., Kathail, P., Carr, A.J., Burdziak, C., Moon, K.R., Chaffer, C.L., Pattabiraman, D., et al. (2018). Recovering gene interactions from Single-Cell data using data diffusion. Cell 174, 716–729.e27. 10.1016/j.cell.2018.05.061.

97. Levitin, H.M., Yuan, J., Cheng, Y.L., Ruiz, F.J., Bush, E.C., Bruce, J.N., Canoll, P., Iavarone, A., Lasorella, A., Blei, D.M., et al. (2019). De novo gene signature identification from single-cell RNA-seq with hierarchical Poisson factorization. Mol. Syst. Biol. 15, e8557. 10.15252/msb.20188557.

